# Leveraging correlations between polygenic risk score predictors to detect heterogeneity in GWAS cohorts

**DOI:** 10.1101/827162

**Authors:** Jie Yuan, Henry Xing, Alexandre Lamy, The Schizophrenia Working Group of the Psychiatric Genomics Consortium, Todd Lencz, Itsik Pe’er

## Abstract

Evidence from both GWAS and clinical observation has suggested that certain psychiatric, metabolic, and autoimmune diseases are heterogeneous, comprising multiple subtypes with distinct genomic etiologies and Polygenic Risk Scores (PRS). However, the presence of subtypes within many phenotypes is frequently unknown. We present CLiP (Correlated Liability Predictors), a method to detect heterogeneity in single GWAS cohorts. CLiP calculates a weighted sum of correlations between SNPs contributing to a PRS on the case/control liability scale. We demonstrate mathematically and through simulation that among i.i.d. homogeneous cases, significant anti-correlations are expected between otherwise independent predictors due to ascertainment on the hidden liability score. In the presence of heterogeneity from distinct etiologies, confounding by covariates, or mislabeling, these correlation patterns are altered predictably. We further extend our method to two additional association study designs: CLiP-X for quantitative predictors in applications such as transcriptome-wide association, and CLiP-Y for quantitative phenotypes, where there is no clear distinction between cases and controls. Through simulations, we demonstrate that CLiP and its extensions reliably distinguish between homogeneous and heterogeneous cohorts when the PRS explains as low as 5% of variance on the liability scale and cohorts comprise 50, 000 − 100, 000 samples, an increasingly practical size for modern GWAS. We apply CLiP to heterogeneity detection in schizophrenia cohorts totaling > 50, 000 cases and controls collected by the Psychiatric Genomics Consortium. We observe significant heterogeneity in mega-analysis of the combined PGC data (p-value 8.54e-4), as well as in individual cohorts meta-analyzed using Fisher’s method (p-value 0.03), based on significantly associated variants.

## 1 Introduction

In recent years Genome-Wide Association Studies (GWAS) have identified thousands of genomic risk factors and generated insights into disease etiologies and potential treatments [1, 2, 3]. Increasingly, there has been interest in advancing beyond these associations towards obtaining a deeper understanding the mechanisms by which genomic factors influence disease [1, 4]. These require models beyond simply combining linear effects of variants, as they often modulate phenotypes indirectly, though the expression of other genes [5, 6].

One such avenue has concerned the apparent heterogeneity of diseases which has not been sufficiently recognized by GWAS: while individuals in cohorts for these studies are frequently classified simply as cases or controls, clinical evidence for several GWAS traits have suggested that there are multiple different subtypes of diseases consisting of distinct sets of symptoms and association with distinct rare risk alleles [7, 8]. For example, polygenic risk scores for major depressive disorder explain more of the phenotypic variance when cases are partitioned into two known subtypes (typical and atypical), and the two subtypes exhibit polygenicity with distinct traits [9]. Similarly, by separating bipolar disorder into its two known subtypes, corresponding to manic and hypomanic episodes, distinct polygenic risk scores comprising different associated SNPs are discovered, with genetic correlation being significantly lower than when individuals are partitioned otherwise, e.g. by batch. Additionally, only the manic subtype shares a high degree of pleiotropy with schizophrenia [10]. Aside from psychiatric traits, heterogeneity of genomic associations between known subtypes has been observed in diseases such as lupus [11], multiple sclerosis [12], epilepsy [13], encephalopathy [14], and juvenile idiopathic arthritis [15]. Elucidating the nature of heterogeneity in these traits may also play a role in addressing the missing heritability problem in GWAS, as hidden heterogeneity reduces power to detect SNP associations [16].

Heterogeneity in disease etiology has also become a concern for clinical applications, as the predictive accuracy polygenic risk scores is known to vary across different demographics of patients. As most genomic studies to date have been conducted on primarily Northern European populations, accuracy of the predictors they develop, measured as R-squared, is lower in other populations, raising the possibility of inequities in care by the direct application of these PRSs [17]. Even if these concerns are mitigated by future large studies conducted in under-served populations, recent work has shown that PRS accuracy further varies across other covariates such as age and sex [18]. Therefore, methods to develop population-differentiated PRSs and detect deficiencies in existing PRSs are urgently needed before predictive genomics can be widely integrated into precision medicine.

To date there have been only few strategies to identify subtypes in GWAS cohorts, largely due to two challenges: the very small signals typically found in polygenic traits, and the presence of confounding sources of heterogeneity such as batch effects. One method [19] purports to discover strong evidence of subtyping in schizophrenia by non-negative matrix factorization of the cohort genotype data, interpreting the hidden factors as different subtypes. However, this work failed to take into account alternative sources of heterogeneity such as population stratification and linkage disequilibrium, that might produce spurious results [20, 21]. Another method, reverse GWAS [22], applies a Bayesian latent factor model to partition SNP effect sizes and individual membership into a set of latent subtypes so that the likelihood of phenotype predictions within each subtype is maximized. The method is reported to detect subtypes that may be suggestive of clinical implications, such as a possible differential effect of statins on blood glucose levels. However, this approach is under-powered to detect heterogeneity in single phenotypes, and thus is geared for simultaneous predictions across multiple observed phenotypes. Additionally, many of these phenotypes are quantitative, which allows for more accurate estimation of effect sizes, and thus more accurate subtyping, than in case/control phenotypes. Therefore methods of this flavor may struggle to detect subtypes among single case/control phenotypes, in which the quantitative liability score is hidden.

Within-phenotype heterogeneity has also surfaced as a possible confounding factor in the discovery of pleiotropic associations between phenotypes [23]. Assuming a GWAS model of disease risk, ideal pleiotropy would involve a single variant significantly associated with two observed phenotypes, producing a genomic correlation between those phenotypes. However, the presence of distinct subtypes in one or both pheno-types may alter the conclusions derived from pleiotropic analysis. For example, two additional subtypes of depression have been characterized by either episodic or persistent experiences of low mood. Of the two, the persistent subtype is more closely associated with childhood maltreatment, and only in persistent cases is an association found between childhood maltreatment and a particular variant of the serotonin transporter gene [24, 25]. Misclassification is another possible source of heterogeneity leading to spurious pleiotropic relationships between phenotypes. For example, a significant percentage of patients diagnosed with either bipolar disorder or schizophrenia have their diagnoses later corrected to reflect the other disease [26]. As bipolar disease and schizophrenia are understood to be highly pleiotropic [27, 28, 29], these misclassifications have the potential to skew analyses of genetic correlation between the two phenotypes.

Recent work by Han et al. [30] has sought to address the detection of heterogeneity specifically in the context of pleiotropic phenotypes. The proposed method, BUHMBOX, operates on a matrix comprising cases for one disease genotyped over the associated SNPs for a second disease. When only a subset of cases are also cases for a second disease, individuals within that subset will exhibit a slightly higher ascertainment for the risk alleles included in the matrix. In a non-heterogeneous pleiotropic scenario, these risk alleles would instead be randomly distributed among all included individuals rather than co-occurring in a subset. When multiple risk alleles are overrepresented in a subset, they are positively correlated across all individuals in the matrix, and these positive correlations serve as evidence of heterogeneity.

We propose a generalized method called CLiP (Correlation of Liability Predictors) that leverages these correlations more broadly to detect multiple forms of heterogeneity in even single-trait GWAS, rather than strictly in two labeled pleiotropic traits. The goals of this work are threefold: First, we demonstrate that in a homogeneous (null) cohort of cases in a case/control study, predictors with effect sizes of the same sign are not uncorrelated as stated by Han et al. [30] but negatively correlated, and are expected to produce negative heterogeneity scores. This is a mathematical consequence of both logistic and liability threshold models despite independent sampling of predictors over the entire cohort. We evaluate the power of CLiP across realistic GWAS scenarios, and demonstrate its utility by identifying heterogeneity in schizophrenia. Although previous methods have attempted to partition SNPs or individuals into distinct clusters, the highly polygenic nature of most phenotypes renders these methods under-powered for single trait GWAS even when data sizes are very large. CLiP aggregates signals across all associated SNPs to generate a single score, permitting users to flag heterogeneous data sets for further study. Second, we develop an extension of CLiP to accommodate parameters that are not binomial genotypes, but rather continuous predictors such as expression data, which we term CLiP-X. Finally, we further extend CLiP to identify heterogeneous subgroups in quantitative phenotypes, where no clear delineation between cases and controls exists, by weighting correlations according to polygenic risk scores, which we term CLiP-Y.

## 2 Methods

From a GWAS perspective, heterogeneity can be interpreted broadly as the presence of distinct mixtures of cases within a cohort which have been identified as cases through different PRSs. We define two models for generating genotype matrices of heterogeneous cohorts: First, *misclassification*, whereby a subset of individuals are not really cases, but have been rather labeled as such despite genetically being controls. This may occur due to erroneous phenotyping, but it may also suggest distinct disease etiologies, some of which are not ascertained for the PRS of interest. Second, a *mixture* of unobserved sub-phenotypes with distinct PRSs. A case is observed if the individual passes the liability threshold of at least one of these sub-phenotype PRSs. Figure 1 displays idealized genotype matrices and correlation matrices for each of these models along with the homogeneous null scenario, in which all cases are selected according to the same PRS. The column set *S* comprises associated SNPs reported in GWAS summary statistics, with the counted allele selected so that the corresponding effect size is positive. As described in Results, associated SNPs participating in the same PRS are negatively correlated over a set of cases selected according to that PRS (panel **B**). When the cohort comprises both cases and misclassified controls, the pattern of ascertainment of risk-alleles is consistent for particular individuals across all SNPs, resulting in positive correlations between SNPs (panel **D**). Panel **E** depicts a mixture scenario with two hidden disjoint PRSs. Individuals labeled as cases of the observed phenotype may be in reality a case for sub-phenotype 1 only (blue), sub-phenotype 2 only (orange), whereas controls are observed as such (grey). The presence of cases for multiple hidden sub-phenotypes produces a mixture of positively and negatively correlated SNPs depending on the membership of the compared SNPs (panel **G**).

**Figure 1:**
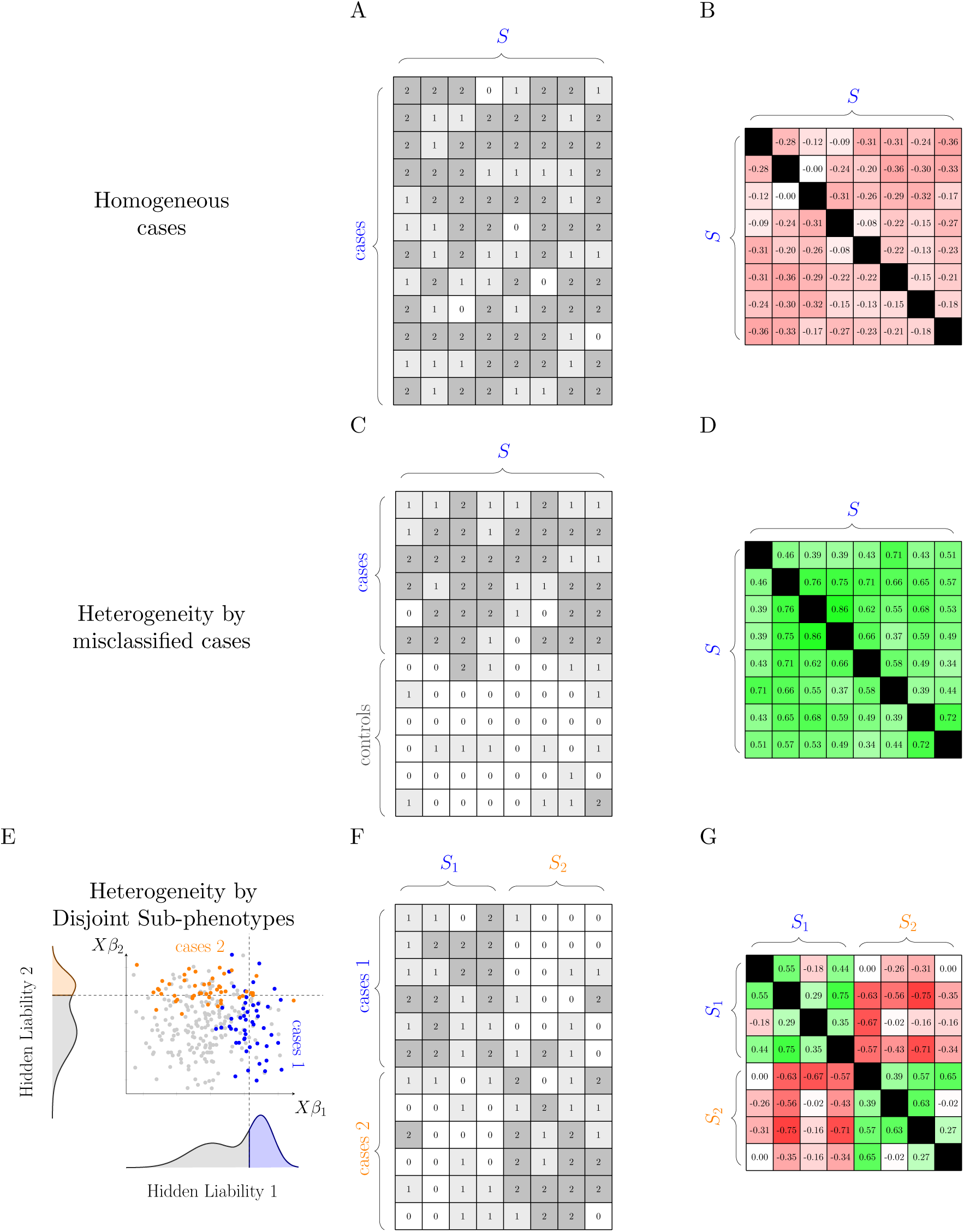
Cartoon examples of genotype matrices **(A,C,F)** and SNP correlation matrices **(B,D,G)** expected of homogeneous and heterogeneous case cohorts. For homogeneous cases **(A,B)**, SNPs are uniformly ascertained, but negative correlations exist between any pair of associated SNPs. For heterogeneous cases comprising a mixture of true cases and misclassified controls **(C,D)**, SNPs are ascertained in a subset of individuals, creating positive correlations between SNPs. For heterogeneous cases comprising disjoint sub-phenotypes **(E,F,G)**, associated SNP subsets *S*_1_ and *S*_2_ pertain to two independent PRSs, and passing the threshold of at least one of these PRSs is sufficient to select a case **(E)**. Genotypes sampled from this model produce a mixture of positive and negative correlations.

The goal of CLiP is to distinguish this heterogeneous cohort from one that comprises only cases and controls for a single PRS. In the following sections, we first describe a correction (CLiP) to the way heterogeneity scores had been used [30], where we account for negative correlations which are expected of case/control data sampled from a logistic or liability threshold model. Next we present adaptations of this general method to studies with quantitative predictors such as expression measurements rather than SNPs (CLiP-X), and also with quantitative phenotypes for which there is no definition of a “case” (CLiP-Y). Next we describe the generative process for simulations of homogeneous and heterogeneous PRS data used to test the performance of these methods.

### 2.1 CLiP: Correcting for negative correlation bias

CLiP calculates the same heterogeneity score as previous work [30], but adjusts the null distribution to account for expected correlations between SNPs when the cohort is homogeneous. Calculation of this adjustment and verification by simulations are shown in the Results and Supplemental Note. The test is performed over a genotype matrix *X* comprising *N* cases and *M* SNPs counting the number of risk-alleles, as well as a matrix of controls *X*^0^ with *N*^0^ individuals. Pairwise SNP correlations are calculated over cases and controls separately and stored in *R* and *R*^0^ respectively. These correlations are then compared against their null expected values. The expected correlation among controls, 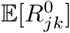, is always 0 in practice as SNPs are sampled independently, but is included below for clarity. This modified heterogeneity score is computed as follows:

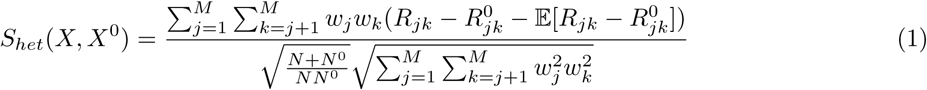

where

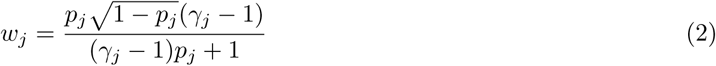

The score *S*_*het*_ is a weighted sum of difference in correlation between cases and controls, to account for prior sources of SNP-SNP correlation such as ancestry. A high score resulting from a bias towards positive correlations would indicate the presence of subtypes with differing ascertainment for the included risk-alleles, and thus heterogeneity. The weights are intended to adjust the score’s sensitivity to certain SNPs based on their allele frequency *p* and odds ratio *γ*, with larger odds ratios and frequencies close to 0.5 producing greater weights.

### 2.2 CLiP-X: Heterogeneity Detection with quantitative predictors

While comorbidity subtypes may occur in transcriptome-wide association studies, the heterogeneity score cannot be computed directly over continuously distributed gene expression variables rather than discrete SNPs. In CLiP, the weights *w* are important for scaling the contributions of individual SNPs to the final heterogeneity Z-score, and they are dependent on risk-allele frequencies and odds ratios, quantities not strictly defined for continuous variables.In the case of binary variables, higher weights are assigned to SNPs with more extreme risk-allele frequencies as well as effect sizes, as these variables are more likely to generate highly positive correlations in the presence of heterogeneity. Here we generalize this weighting scheme to accommodate arbitrarily distributed continuous input variables, which may be applied in particular to expression analyses.

### 2.3 CLiP-X Simulation Procedure

To fully simulate expression variables as modeled in transcriptome-wide association, expression predictors are generated from a linear model of randomly sampled genotypes, rather than directly sampling expression. Although the input into CLiP-X includes only the expression variables, explicitly modeling the genotype layer allows for inclusion of prior correlations resulting from SNPs associated with multiple transcripts, rather than from the liability threshold model.

For a single case-control phenotype, transcript effect sizes *α* are fixed so that the variance explained of all modeled transcripts is a desired value. Likewise, genotype-transcript effect sizes *β* are also fixed so that variance explained of each transcript by genomic variants is a second specified value. Although fixing effect sizes at the genotype-transcript layer is admittedly unrealistic, the results are only simplified when these interactions are removed, with no interactions reducing to expression sampled from the standard normal distribution. Cases are determined according to the liability threshold model. For an individual *i* in transcript matrix *Z*, a hidden quantitative liability score 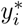 is calculated, with the variance of error *ϵ* set so that *y*\* has a total variance of 1. The observed case/control label *y*_*i*_ is set according to whether 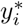 passes the liability scale threshold *T*, which is placed on the standard normal distribution so that affected individuals constitute a prevalence of 0.01.

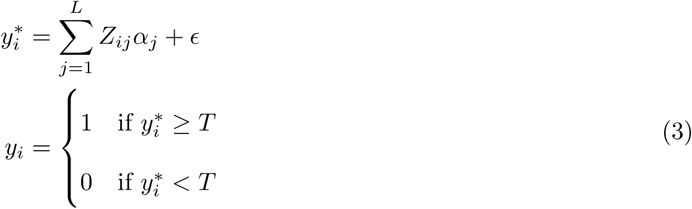

To generate cases and controls, we iteratively generate batches of transcripts by random sampling, and compile those that pass or fail the threshold cutoff into case and control cohorts. We generate heterogeneous cohorts, by concatenating simulated cases and controls, with the fraction of cases *π* set to 0.5 for simplicity. We reasoned that this procedure was a conservative representation of a large number of possible heterogeneity scenarios including those with multiple independent sub-phenotypes. If the PRSs of these sub-phenotypes are independent, then a large number of correlations between predictors will be evaluated close to zero, resulting in a score very different from the homogeneous null. A full description of the simulation procedure is provided in Supplemental Algorithm 1 and illustrated in Supplemental Figure S2. Note that the variance of the random noise *ϵ* in Equation 3 is determined by the desired total variance explained by the simulated genotypic variables *h*^2^:

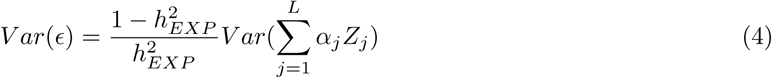

#### 2.3.1 Characterizing correlations between continuous variables

Given *N × L* matrices of quantitative expression measurements *Z* among cases and *Z*^0^ among controls, we would like to determine whether *Z* comprises a homogeneous or heterogeneous set of cases as generated in Supplemental Algorithm 1. When *Z* is heterogeneous, we assume the individuals in *Z* can be assigned to one of two subtypes: one sampled according to the liability threshold model for the simulated phenotype, and one sampled randomly as controls. For a given predictor indexed by *j* ∈ [1, …, *L*], assume *Z*_*ij*_ is sampled according to a mean and variance specific to the subtype of individual *i*, denoted by 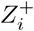 for the case subtype and 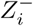 for the control subtype. The distribution of the variables need not be discrete or even normally distributed, as the heterogeneity score is computed from correlations, which in turn rely only on the mean and variance of the input variables. Therefore the score can be calculated assuming any probability distribution provided that the mean and standard deviation are obtainable. For an arbitrary probability distribution *𝓓* parameterized by its mean and standard deviation, we have:

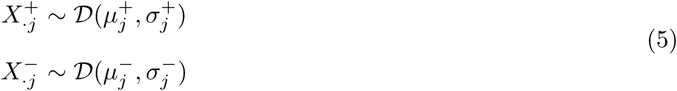

Assume that the proportion of individuals belonging to the group + is *π*. For a homogeneous group of case, *π* = 0, and our simulations assume *π* = 0.5, but in practice this proportion is unknown. Incorporating this proportion allows the redefinition of expectations over the entire cohort as weighted sums of the expectations over the subgroups. The expected correlation evaluated over the entire group can then be calculated according to within-group expectations:

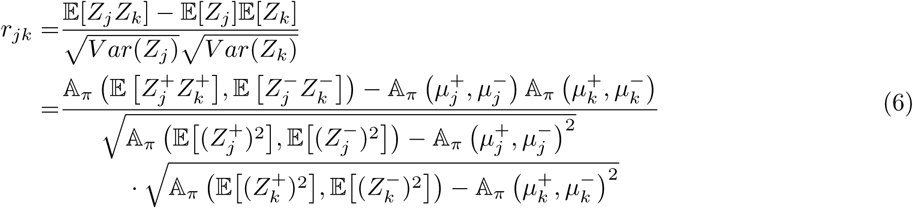

where 𝔸_*π*_(*x, y*) = *πx* + (1 − *π*)*y*.

#### 2.3.2 Definition of weights for continuous variables

We would like to make use of these expectations over correlations by incorporating them as weights in the heterogeneity score as in Han et al. [30]. As predictors with high mean differences between subgroups and high effects are expected to contribute more signal to the score, weighting them higher than other predictors will increase power to detect heterogeneity. Therefore, we would like to define a set of weights *w*_*ij*_ for each expected *r*_*ij*_.

We derive the weights for continuous variables in an analogous manner to Han et al. [30], by taking the derivative of the expected sample correlation with respect to *π* at the null value, *π* = 0.

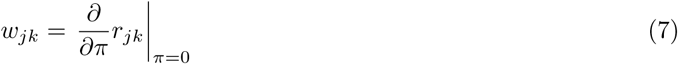

To facilitate calculation of 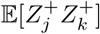 and 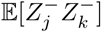 in equation 6, we assume as in [30] that within a subgroup of cases or controls, the correlation between any two predictors, even those associated with the phenotype, is zero. This allows us to express expectations of products as products of expectations. Note that this does not mean that correlations over the entire cohort 𝔼[*Z*_*j*_*Z*_*k*_] are zero: these correlations are calculated inclusive of all subgroups, and their nonzero correlations are what determines the heterogeneity score. We demonstrate in the Results that theoretically and by simulation this assumption is violated in logistic and liability threshold models.

Given the assumption of no correlation within subgroups, the correlation between two variables *Z*_·*j*_ and *Z*_·*k*_ can be expressed as the following. For further details on the derivation, please see the Supplemental Note.

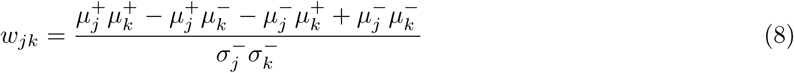

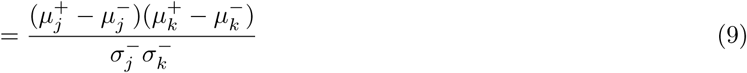

The same weights defined in Han et al. [30] for Bernoulli variables is a special case of this general formulation. These weights can now be substituted into the heterogeneity score.

In practice we do not know the value of 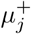 because the membership of individuals in each of the subsets is unknown. However, we do know the mean values of the heterogeneous case group which we denote as *µ*_*j*_. We can use this value as an approximation for 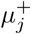, and calculate an approximate weight:

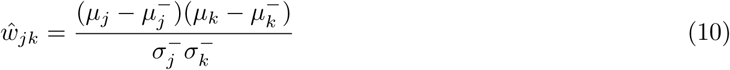

We can also quantify the errors we are making by this approximation. We have the following relationship for any distribution of the genotype random variables:

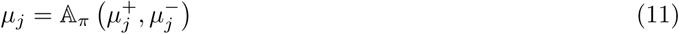

The approximation in Eq. 10 will attenuate the magnitude of 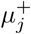 with respect to the true value of the weight. However, we also see that:

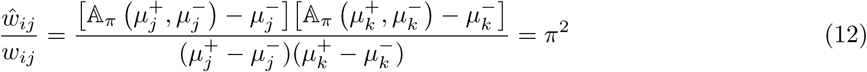

As each weight is scaled by a constant factor, their relative magnitudes are unchanged. Consequently, the heterogeneity score for continuous input variables does not change after this approximation. Thus we can still achieve optimal estimates of heterogeneity despite lacking access to the true mean for the underlying case subgroup.

### 2.4 CLiP-Y: Heterogeneity Detection in Quantitative Phenotypes

The basic CLiP test for heterogeneity relies on differential enrichment of SNP effect sizes or odds ratios across subtypes, and thus requires ascertainment for cases. But one can presume that heterogeneity exists in quantitative phenotypes as well; e.g., are there distinct genetic mechanisms predisposing individuals to being tall? But extending this method to quantitative phenotypes presents a challenge as there is no dichotomous delineation between cases and controls. A naive solution may be to pick an arbitrary z-score as a threshold and denote samples who score higher as “cases” and those lower as “controls.” This introduces a trade-off between sample size and signal specificity, as lowering this threshold provides more samples for the correlation analysis but also introduces more control-like samples which will attenuate SNP associations, and the correlations themselves. A more principled method would allow for the inclusion of all continuous samples, but give higher weight to those with large polygenic SNP scores. Thus we propose to score heterogeneity by a weighted correlation with polygenic risk scores serving as a measure of the importance of a sample in the case set. These weights determine the degree to which individuals count as a “case”, and therefore their contribution to the total heterogeneity score of the genotype matrix. Artificially creating the two groups by applying a hard threshold over the quantitative phenotype values is a special case of this method with a step function as the weighting scheme, equally weighting all individuals above the threshold “step.”

### 2.5 CLiP-Y Simulation Procedure

Here SNPs as input predictors are sampled directly from binomial distributions with fixed minor allele frequency of 0.5. The quantitative phenotype *y* is calculated from the PRS score with normally distributed noise added according to the desired PRS variance explained. As in the CLiP-X simulation procedure, we generate heterogeneous cohorts by concatenating a subset of cases and controls together into a single putative set of cases according to the fraction *π*. For quantitative phenotypes, the “control” subset is generated so that the quantitative phenotype value is simply sampled from the normal distribution with zero PRS variance explained. A more detailed description of the simulation procedure is provided in Supplemental Algortithm 2.

#### 2.5.1 Definition of individual weights by phenotype values

We define a weight over individuals such that those with higher phenotype values contribute more strongly to the heterogeneity score. For a cohort of *N* individuals let *X*_*ij*_ ∈ {0, 1, 2} be the number of risk alleles of SNP *j* in individual *i*, and let *Y* = (*y*_1_, …, *y*_*N*_) values of quantitative trait 1 for the respective individuals. We introduce a normalized weight vector across the *N* individuals defined as ***φ*** ∈ ℝ^*N*^ such that *∀i, φ*_*i*_ ≥ 0 and 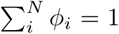. Most intuitively we would define ***φ** ≡ **φ***(*ℱ*), where the weight values would reflect normalized scaling of the trait 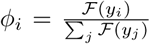 by a monotone function *ℱ*. Dichotomous, case/control weighting is the special case of:

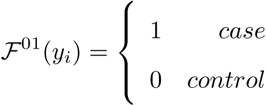

Uniform weighting is obtained by *ℱ*^1^(*y*_*i*_) ≡ 1. To obtain the optimal weight function which most clearly contrasts the scores of heterogeneous and homogeneous cohorts, we tested several possible functions and also performed a local search over polynomials of arbitrary degree by iteratively updating and testing the performance of individual polynomial coefficients. This local search is described in detail in Supplemental Algorithm 3. First, a small number of homogeneous and heterogeneous cohorts are generated as described before. These serve as the training data by which the weight function is optimized. All weight functions are applied over the raw phenotype values directly, or their conversion to percentiles in the sample distribution, in the range [0,1]. After initially randomizing a set of coefficients, at each iteration, a coefficient is randomly selected and incremented by a random quantity sampled from a normal distribution. The resulting polynomial is tested against the training data, and the change to the coefficient is kept if the difference in score between heterogeneous and homogeneous cohorts increases. After a set of high-performing weight functions are selected, they are each evaluated against a larger sample of validation data comprising homogeneous and heterogeneous cohorts as before. Of these candidates, the polynomial that performs best on the validation data is selected.

#### 2.5.2 Definition of weighted correlations

To compute correlations we define, for each SNP *j*, a random variable 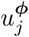 with values in {0, 1, 2} by sampling from the genotypes of the sample cohort *X*_·*j*_ with probability equal to the weight *φ*_*i*_ assigned to each individual *i*. Rather than calculate the correlations directly over SNPs in *X*, we now calculate correlations over these random variables. We omit the superscript ***φ*** in *u*^*φ*^ when it is clear from context. For a single SNP *j*, we define the weighted mean value across *N* individuals as:

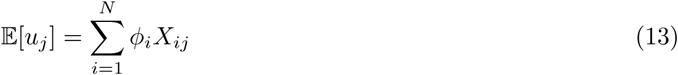

Between two SNPs *j* and *k*, we define the weighted covariance as:

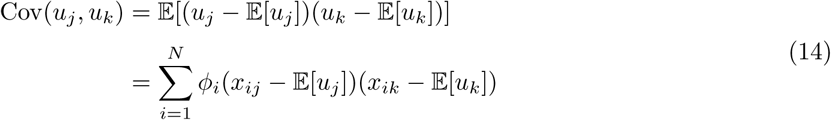

We define the weighted correlation matrix *R*^***φ***^ for any weighting ***φ*** as:

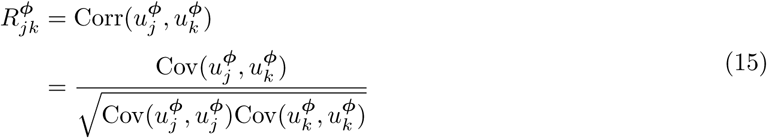

The heterogeneity score tallies the entries of the upper-triangular correlation matrix for the phenotype-weighted individuals *R*^***φ***(*ℱ*)^. As we now lack a held-out set of controls to cancel the contribution of correlations unrelated to the phenotype, we instead calculate a conventional correlation uniformly weighted across all individuals 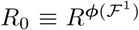. Additionally, we introduce a scaling factor of 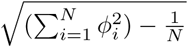 to correct for the change in variance resulting from re-weighting the correlation according to individual weights *φ*_*i*_. These changes produce the following preliminary heterogeneity score for quantitative phenotypes:

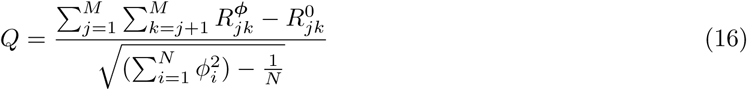

Lastly, we incorporate into the test statistic *Q* a weighting scheme over SNPs as described in Han et al. [30]. This second set of weights ***w*** ∈ ℝ^*M*^ is introduced to correct for larger contributions to the score by SNPs with large effect sizes or risk allele frequencies close to 0.5. These weights apply to SNPs, and should not be confused with the weights ***φ*** over individuals. For each SNP *j*, we define 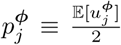, the sample allele frequency weighted by the individual phenotype, as opposed to the unweighted allele frequency 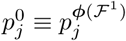. The contribution of SNP *j* to the heterogeneity score is then scaled by

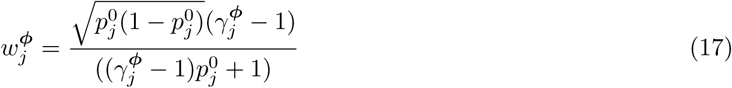

where

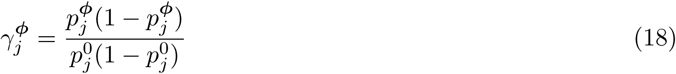

is a weighted generalization of an odds ratio. These weights are analogous to those found in Han et al. [30], where given case allele frequency 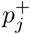, control allele frequency 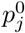, and sample odds ratio 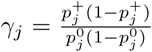, the weight is

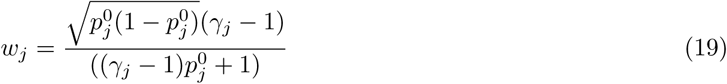

Combining these intermediate calculations, the heterogeneity test statistic for continuous phenotypes is:

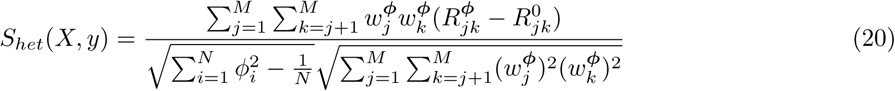

For high *N*, this test statistic approaches the standard normal distribution, and can be evaluated as a z-score hypothesis test.

Note that even when applying a dichotomous weighting scheme, dividing the cohort with quantitative phenotypes into artificial cases and controls, CLiP-Y still differs slightly from a direct application of the case/control score. If a dichotomous weight function produces *N*^*φ*^ artificial cases, the scaling factor 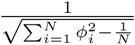 simplifies to 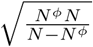 instead of the slightly smaller 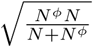 in the original case/control score. This corrects for the slight reduction in variance of 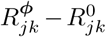 because these differently-weighted correlations are taken over a single cohort of individuals rather than disjoint sets of cases and controls. In practice, we find this correction factor performs very well in scaling the test statistic variance to 1.

### 2.6 Evaluating heterogeneity in SCZ

We applied CLiP to test for heterogeneity in case/control data for schizophrenia collected by the Psychiatric Genomics Consortium (PGC). The data comprise in total roughly 23,000 cases and 28,000 controls and was the subject of a 2014 meta-analysis reporting 108 schizophrenia-associated loci [31]. We would like to test whether heterogeneity suggested from clinical observation is also detectable at the level of the PRS comprising these loci. The PGC data is an aggregate of cohorts collected from many studies conducted in different populations. Therefore a test for heterogeneity over the all cohorts is likely to be confounded by ancestry stratification or batch effects between cohorts. We attempt to circumvent these confounding variables by applying GWAS meta-analysis methods to CLiP scores evaluated over individual cohorts, as well as evaluating the p-value of the sum of all CLiP scores. As each CLiP score is standard normal distributed over the null, the distribution of their sum has expectation 0 and standard deviation 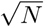 if *N* is the number of cohorts in the sum. To evaluate the significance of CLiP Z-scores across individual cohorts, we applied Fisher’s method for summing p-values [32].

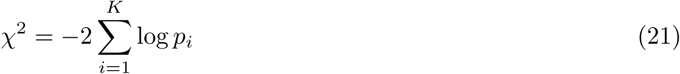

where *K* is the total number of cohorts and *p*_*i*_ is the p-value of the CLiP heterogeneity score for cohort *i*. The p-value of this test statistic is evaluated on a chi-square distribution with 2*K* degrees of freedom. Additionally, we calculated the meta-analysis Z-score of the CLiP score in a manner analogous to the conventional GWAS approach, but with a 1-tail test for highly positive scores only. The meta-analysis Z score is calculated according to

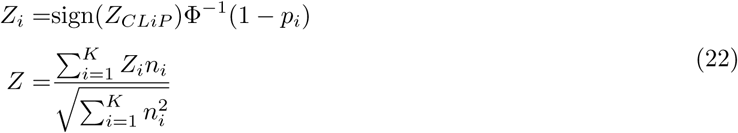

where *Z*_*CLiP*_ is the CLiP Z-score evaluated against the expected score with a standard deviation of 1, and *n*_*i*_ is the sample size of cohort *i*. The results of individual cohort tests along with meta-analysis tests are shown in Table 1.

**Table 1:**
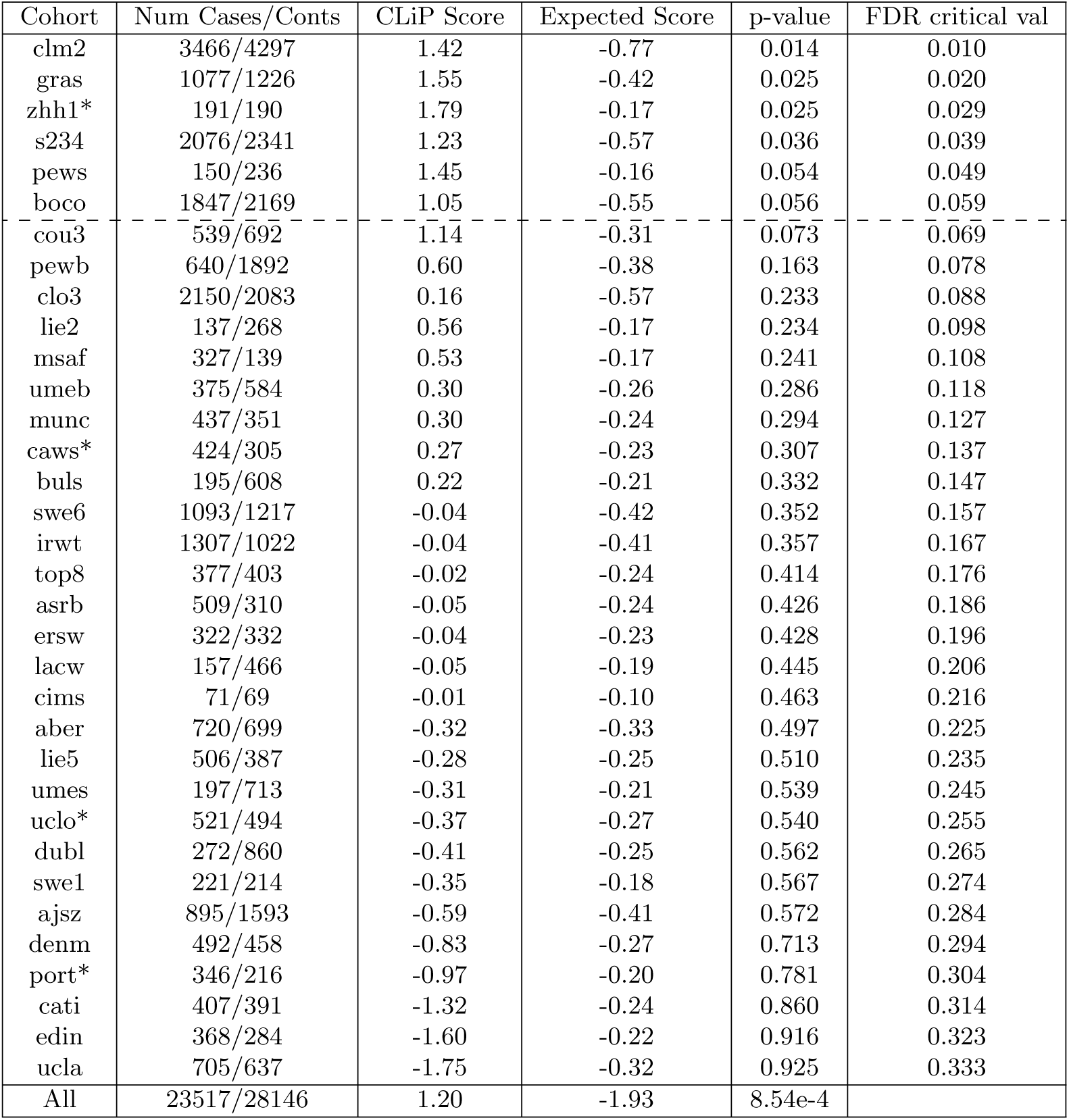
CLiP heterogeneity scores for individual cohorts and their combination. Cohorts with an asterisk have had a SNP excluded which has zero variance within either the case or control cohort, resulting in an undefined correlation. An FDR of 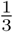 was used for Benjamini-Hochberg analysis.

### 2.7 Application to GWAS of Schizophrenia

We applied CLiP to GWAS data from the PGC, phased and imputed using SHAPEIT [33] and IMPUTE2 [34], a pipeline with similar or better accuracy compared to other tools according to a recent evaluation [35]. Imputation was performed using the 1000Genomes Phase 3 reference panel. Roughly half of the PGC cohorts were mapped with assembly NCBI36, and the SNP coordinates of these data sets were converted to GRCh37 using the LiftOver tool in the UCSC genome browser database [36]. Individuals were excluded from further analysis if their percentage of missing data was greater than 0.1 in the 1 Mb region flanking each SNP. Additionally, of the 108 associated SNPs and indels reported in Ripke et al. [31], six were excluded because they are not listed in the 1000Genomes Phase 3 reference panel, one was excluded due to low variance in many individual study cohorts, and one was excluded due to mismatching alleles between reported summary statistics and the reference panel, for a total of 100 variants included in the heterogeneity analysis.

To accurately estimate expected heterogeneity scores, the odds ratios reported in Ripke et al. [31] must be converted to effect sizes on the liability scale. We apply an approximate method reported by Gillett et al. [37] to convert for variant *j* an odds ratio *OR*_*j*_ to the liability effect *β*_*j*_:

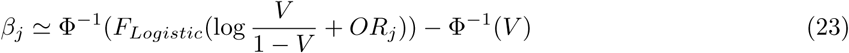

where V is the disease prevalence (0.01 for schizophrenia), and 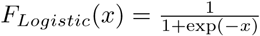.

## 3 Results

### 3.1 Implementation of CLiP

We have implemented the CLiP family of methods with open source availability of the software and auxiliary code for generating results reported in this paper, available at https://github.com/jyuan1322/CLiP. For the scenarios reported in this manuscript, the runtime to simulate and test a single cohort is always below 5 minutes on a standard machine.

#### 3.1.1 Correlations between effect variables in cases

One of the fundamental claims of Han et al. [30] is that SNPs conferring risk for a disease are uncorrelated among cases for the disease as well as controls. However, the authors prove this using only for a multiplicative binary model, in which an individual’s risk is the product of odds ratios of probability of disease for associated SNPs. The most common model in contrast is a logistic or liability threshold model, in which these odds ratios are thresholded by a sigmoid function, potentially introducing correlations between SNPs. In practice the results of logistic or liability threshold regression are very similar, with effect sizes differing by approximately a constant factor [37]. We first tested the conventional, case-control score for heterogeneity, as implemented in Han et al. [30], with cases generated from a full logistic model as well as the multiplicative model. We simulated individual data using 100 independent SNPs whose effects were standard-normally distributed. Across logistic and multiplicative models, the same odds ratios are ascribed to each SNP to facilitate comparison. Also, for simplicity, we assumed a variance explained of 1 to better observe the resulting correlation signal. We evaluated the dichotomous-trait score relative to sample sizes for both homogeneous and heterogeneous groups. The result is shown in Figure 2A, where it is apparent that the behavior of the two models diverges drastically. In particular, when a logistic model is assumed the null test statistic is significantly biased towards negative values, indicating widespread negative correlations between the SNPs that contributed to the liability score, in contrast to the multiplicative model considered by Han et al. [30]. While true heterogeneity results in positive scores as before, in the logistic model these scores are also highly attenuated by the negative bias observed in controls. A more detailed discussion of these negatively correlated SNPs can be found in the Supplemental Material.

**Figure 2:**
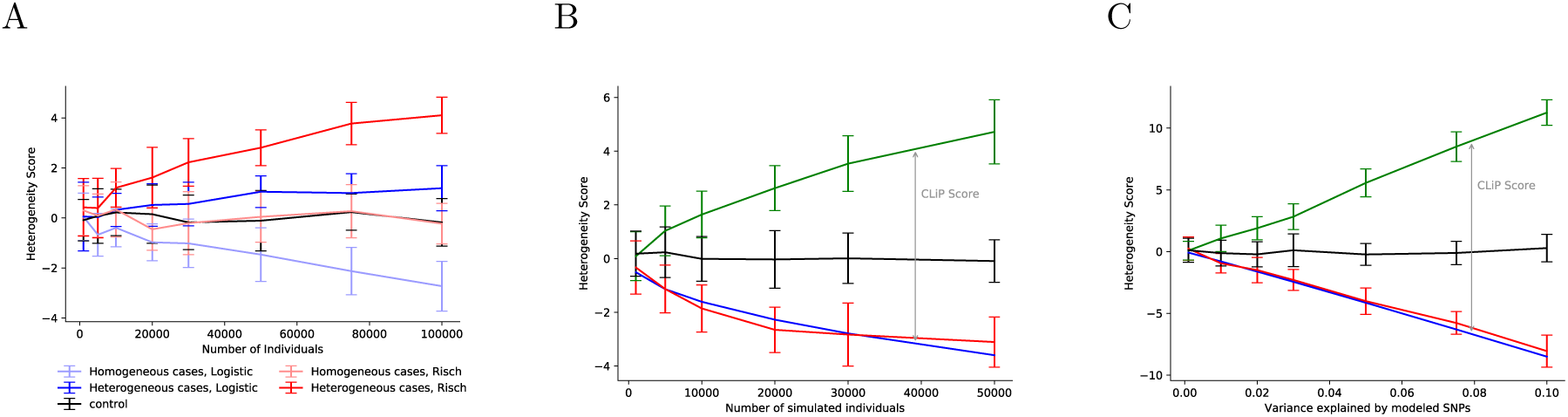
**(A)** Heterogeneity scores (y-axis) across different cohort sizes (x-axis) and genetic architectures. Thresholded models for case/control data such as logistic and probit (liability threshold) regression produce negative correlations between predictors, while the simpler multiplicative (Risch) model does not. Here case/control cohorts are generated from logistic or Risch models with 10 diploid SNPs with allele frequency 0.5 and OR 1.16 (set to keep Risch probabilities ≤ 1). As described in [30], Risch model cases exhibit no correlations in homogeneous cases, and positive correlations in heterogeneous cases, producing zero and positive heterogeneity scores, respectively. However, in thresholded models, negative correlations in homogeneous models produce negative scores. This negative bias in homogeneous scores is unaccounted for in the method by [30], significantly increasing the probability of type II errors. **(B)** Heterogeneity scores (y-axis) on simulated case/control cohorts as a function of sample size (x-axis) with a fixed variance explained of 0.034 as in [31]. Simulations are run with a PRS of 100 SNPs with total variance explained of 0.034. Heterogeneous cohorts (Green) are equal-proportion mixtures of controls (Black) and homogeneous cases (Red). The expected homogeneous score (Blue) is calculated from effect sizes and allele frequencies of PRS SNPs only, and should be used as the true null score in CLiP. **(C)** Heterogeneity scores (y-axis) as a function of variance explained (x-axis) with a fixed sample size of 30,000 cases and 30,000 controls.

### 3.2 Correction for Negative Correlation Bias

To demonstrate the effects of correlated predictors on heterogeneity detection, we evaluated hetrogeneity scores on simulated homogeneous and heterogeneous cohorts. Simulation parameters were set to approximate those described in a meta-analysis of schizophrenia GWAS by the Psychiatric Genomics Consortium [31], which describes a PRS over 108 genomewide-significant SNPs with a total variance explained of 0.034, typical of current GWAS for highly polygenic phenotypes. Genotypes over 100 associated SNPs are sampled according to a fixed risk-allele frequency of *p* = 0.2. Effect sizes are set to a fixed value producing the desired variance explained in a standard normal PRS distribution. Homogeneous case cohorts were generated by repeatedly sampling control genotypes and selecting individuals whose PRS pass a threshold corresponding to a prevalence of 0.01. Heterogeneous cohorts are created by combining an equal number of homogeneous cases and controls. The scores of these cohorts were evaluated over a range of sample sizes keeping variance explained constant at 0.034 (Figure 2B), and a range of total variance explained keeping sample size constant at 30,000 cases and 30,000 controls (Figure 2C). Additionally, we tested the performance of CLiP with respect to the fraction of individuals in the case mixture that are true cases, shown in in Figure 3A. The color of each line indicates the size of the entire case cohort, while the X-axis indicates the fraction of individuals of that count that are true cases. When the fraction is 0, the cohort contains only controls, and all expected correlations are 0, producing a heterogeneity score of 0. When the fraction is 1, the cohort contains only cases, and produces a highly negative score due to negative correlations between all pairs of SNPs. As expected, a mixture of cases and controls produces positive scores, with the peak score occurring when the cohort is split evenly. More detailed results of this set of simulations are shown in Supplemental Table S2.

**Figure 3:**
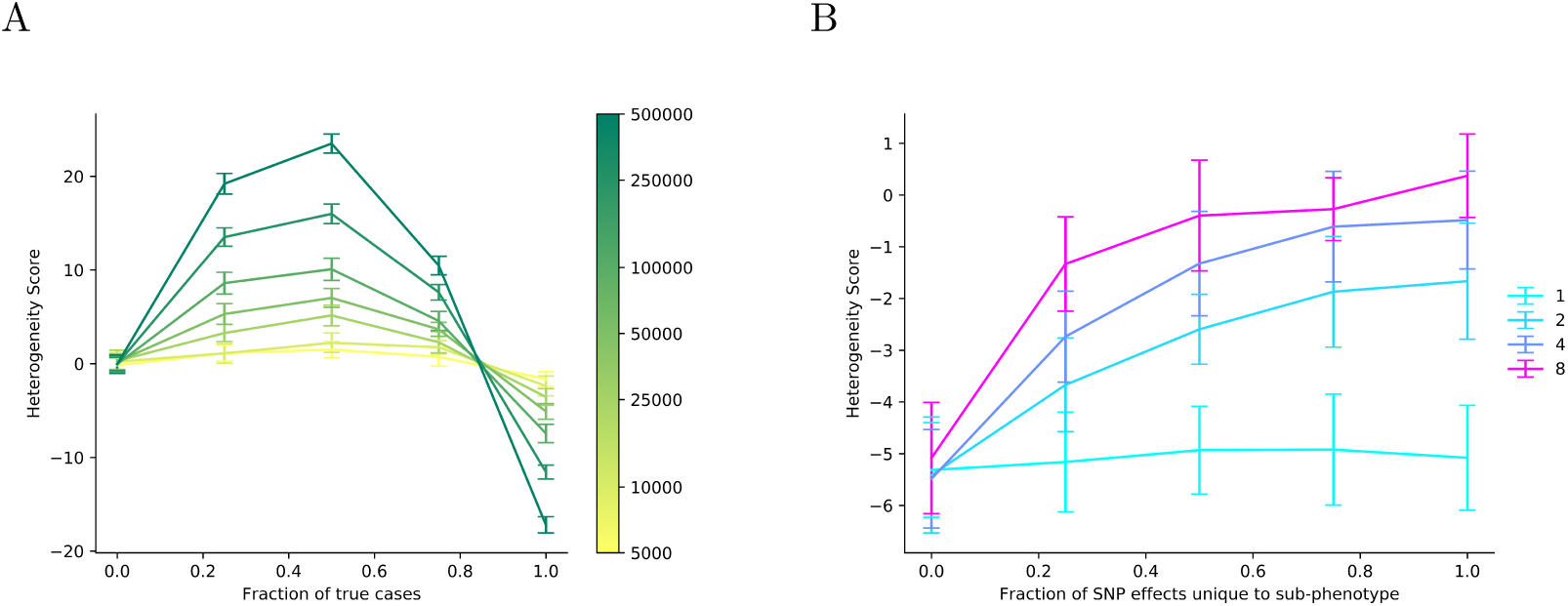
**A** Heterogeneity scores (y-axis) evaluated on heterogeneous cohorts comprising a mixture of true cases and controls at different proportions (x-axis). Colors indicate the total cohort size. All tests were conducted over 50,000 cases and 50,000 controls, with a SNP variance explained of 0.05. **B** Heterogeneity scores (y-axis) evaluated on simulated heterogeneous cohorts with disjoint sub-phenotypes. Performance is shown a function of the fraction of SNP effects unique to a particular sub-phenotype (x-axis). Colors indicate the number of sub-phenotypes. Simulations were performed with 50,000 simulated cases and 50,000 controls, and a total SNP variance explained over all sub-phenotype PRSs set to 0.05.

We also evaluated the performance of CLiP when heterogeneity consists of multiple potentially independent sub-phenotypes, each with a distinct PRS, such that an individual is considered to be a case when it is a case for one or more of these sub-phenotypes. Discovering heterogeneity in these cohorts is more challenging because correlations between SNPs involved in different sub-phenotype PRSs are expected to be zero rather than positive, and if there are any SNP associations shared between sub-phenotypes, negative correlations will be expected between them despite the presence of distinct sub-phenotypes. We tested the performance of CLiP by fixing the number of cases and controls at 50,000 each, the total number of SNPs at 100, and the total variance explained at 0.05, while varying the number of sub-phenotypes and the fraction of SNPs that are shared across all sub-phenotypes. When this fraction is zero, the sub-phenotypes are completely independent, and the SNPs are divided into mutually exclusive subsets associated with each sub-phenotype. When the fraction is non-zero, that fraction of SNPs has the same effect size across all sub-phenotypes. Results of these simulations are shown in Figure 3B as well as Supplemental Table S1. Note that by dividing associated SNPs into associations with particular sub-phenotypes, the total variance explained for each sub-phenotype is reduced, and the observed variance explained of the entire heterogeneous cohort will be lower in a simple linear regression.

### 3.3 CLiP-Y: Quantitative Phenotypes

In practice, we found converting PRSs to percentiles improved performance for all learned weight functions, possibly because percentiles limit the domain of the PRS function over which the function must be *≥* 0, and they reduce the contribution to the score calculation by extreme PRS values. We performed this search for polynomials of increasing degree, finding optimal polynomial functions show in Figure 4A. All polynomial functions converged to highly similar concave functions. This is due to the balancing effect of the normalization factor on the sum of correlations: while correlations of PRSs at the high end of the distribution are more extreme because these individuals more closely resemble “cases,” a high weight value at the higher end of the PRS spectrum means that the normalization factor also shrinks the magnitude of the score. To demonstrate that optimal weight functions are concave functions over the range of PRS percentiles, we tested weight functions that sum up two indicator functions for intervals in [0, 1], one increasing, for an interval ending at 1, and another decreasing, for an interval starting at 0, and evaluated heterogeneity detection performance, shown in Figure S9. The best performing functions are those where the increasing function threshold is near but not at 0, and the decreasing function threshold is near but not at 1, producing a function similar to the concave polynomials found in Figure 4A.

**Figure 4:**
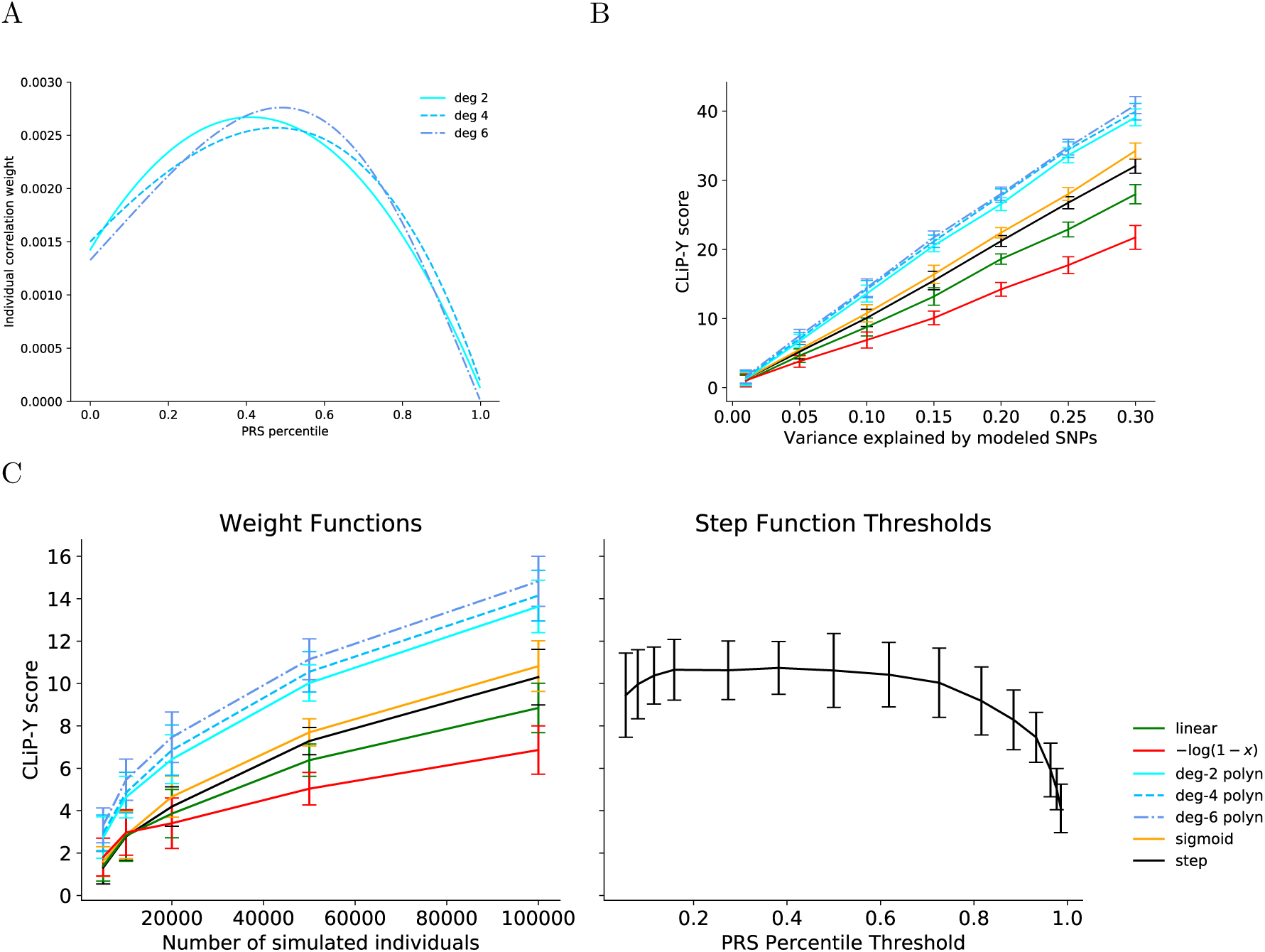
**A** Learned weight functions *φ*(*x*) for scoring heterogeneity in quantitative phenotypes. A local search over polynomial coefficients is performed such that the resulting function maximizes the difference between the heterogeneity scores of simulated samples of homogeneous and heterogeneous cohorts. **B** Tests for heterogeneity in quantitative phenotypes using multiple weighting functions over individuals, including those in **A**, as a function of variance explained by the PRS. Plotted are scores of heterogeneous cohorts minus the expected score of a homogeneous cohort over 20 trials. One Hundred SNPs are simulated with cohorts of 100,000 individuals. **C** Tests for heterogeneity in quantitative phenotypes using multiple weight functions. Plotted are mean scores of heterogeneous cohorts minus the expected score of a homogeneous cohort over 20 trials, as a function of sample size. One hundred SNPs are simulated with a total variance explained of 0.1. For comparison these scores are plotted on the same Y-axis as scores generated from step function weights at various thresholds on the percentile scale of a standard normal quantitative phenotype distribution. For each of these step function scores, the expected homogeneous score is estimated by the mean of 20 sampled homogeneous cohorts, to limit computation time. For all weight functions and test conditions, expected homogeneous scores are near-exact estimates of the means of simulated scores, as shown in Supplemental Figures S5A and S7A.

In the absence of a method for scoring continuous phenotypes, a naive approach using conventional heterogeneity scoring [30] would involve setting an arbitrary PRS cutoff by which to partition the cohort from a continuous phenotype into cases and controls. We compare our continuous heterogeneity test to cutoffs at various percentiles of PRS’s. Both the continuous heterogeneity test and the arbitrary cutoff tests are standard normally distributed in the null scenario, when no heterogeneity is present. As shown in Figure 4, the continuous heterogeneity test outperforms all thresholded tests by achieving the highest score when heterogeneity is present. This is consistent across all tested simulation parameters for total genomic variance explained and number of individuals in the entire cohort. Also note that as expected, the best performing z-score threshold is some intermediate rather than extreme value. Reducing the threshold too much adds to much noise to the correlation, and conversely raising it too high reduces the number of cases and hence the detectable signal too much. From the tests shown in Figure 4, a threshold near the center of the PRS distribution seems close to optimal, but this is significantly outperformed by the continuous method. Plotted here are the differences in heterogeneous and homogeneous cohorts, with the homogeneous cohorts being the true null value of the score. The scores for these cohorts individually are shown in Supplemental Figures S4 to S7.

### 3.4 CLiP-X: Quantitative Predictors

We generate cases and controls for both homogeneous and heterogeneous transcriptome-wide association cohorts, with 100 simulated SNPs generating 10 transcriptome-level variables. We run 20 trials across a range of sample sizes and total genomic variance explained, controlled by the value of *ϵ* in equation 4, to evaluate the performance of the continuous variable test statistic in true heterogeneous cohorts, homogeneous case cohorts, and independently sampled control cohorts. The results are presented in Figure 5 and Supplemental Figure S3.

**Figure 5:**
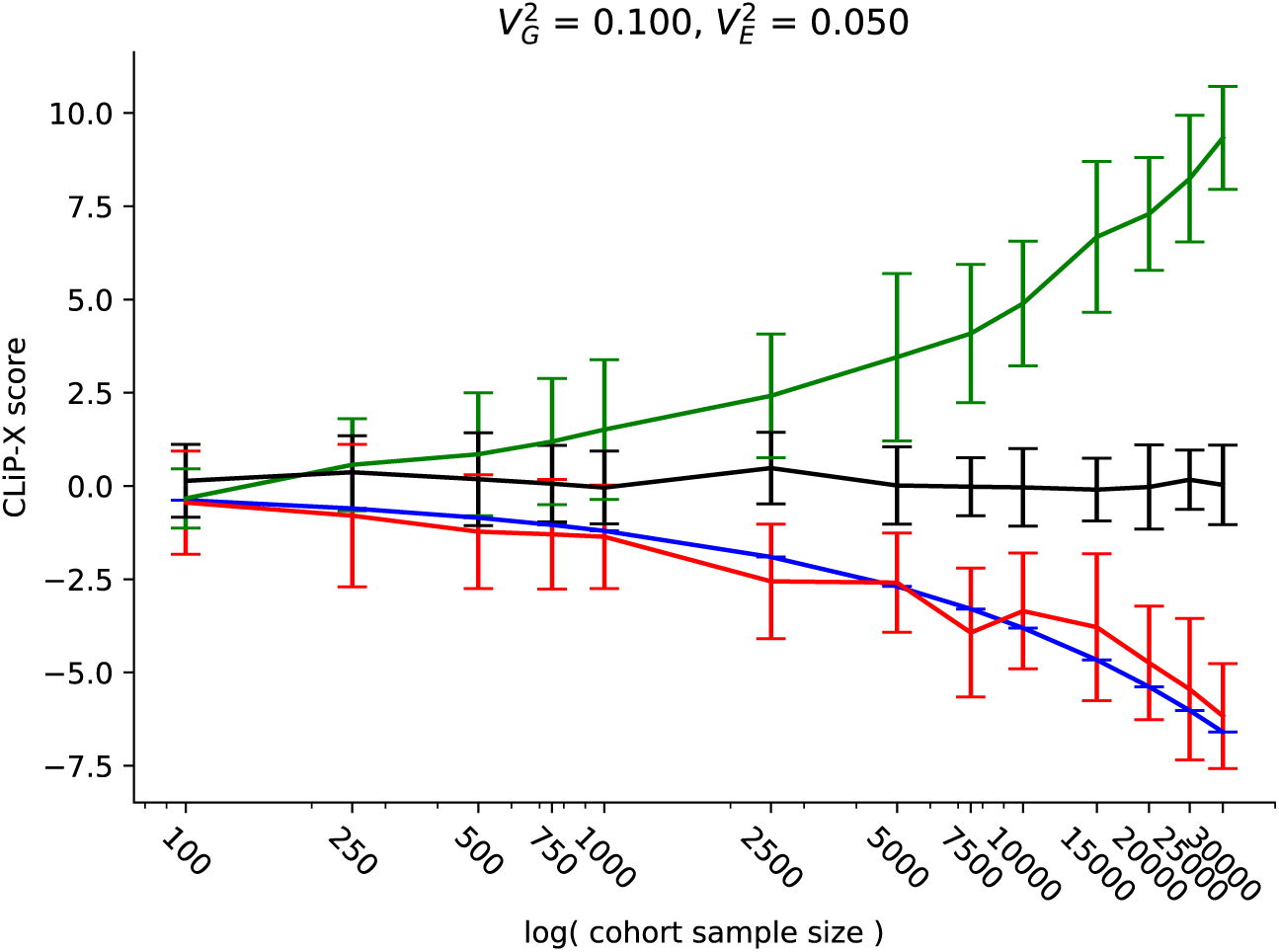
Heterogeneity scores with continuous input predictors generated according to Supplemental Algorithm 1 and Supplemental Figure SS2. Controls **(Black)** have no criteria for selection placed on their generated quantitative predictors; homogeneous cases **(Red)** are selected according to a liability threshold over predictors; and heterogeneous cases **(Green)** are an even combination of controls and homogeneous cases. The **Blue** line indicates expected mean scores of homogeneous cohorts calculated from summary statistics of the quantitative predictors. As with discrete SNPs, quantitative predictors are negatively correlated among homogeneous cases.

Shown in Figure 5 are results with a sample size of 100,000 and a total variance explained of 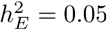 by quantitative predictors. We observe that for all sample sizes, the heterogeneity score is approximately distributed with mean 0 and standard deviation 1 in control cohorts. As predicted, the homogeneous case group exhibits highly negative correlations between associated SNPs, and the resulting CLiP-X score can be accurately estimated from expected correlations (Blue) using knowledge of summary statistics only. This estimate should serve as the null when evaluating GWAS cohorts in practice, when a truly homogeneous cohort is not available. By comparison to this true null, many more heterogeneous cohorts are detectable which would not have passed a significance threshold with the null centered at 0, especially those with sample sizes of less than 10,000 cases.

### 3.5 Application to GWAS of Schizophrenia

After transforming PGC effect sizes to the lability scale (see Methods), the total variance explained by the 100 genomewide significant SNPs considered (see Methods) was approximately 0.027, suitably close to the 0.03 SNP variance explained reported in Ripke et al. [31]. We calculated heterogeneity scores for cases and controls over individual cohorts, shown in Figure 6, as well as meta-analysis scores over all cohorts as described in the methods, shown in Table 1. Generally, we observe more positive heterogeneity scores for larger cohorts, though only three pass a significance p-value threshold of 0.05. The scores in Table 1 are organized by ascending p-value, and a Benjamini-Hochberg procedure is conducted with a false-discovery rate of 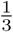. Cohorts with p-values lower than the critical values determined by this FDR are separated by a dashed line. On an individual basis the vast majority of these cohorts are too small to be conclusively tested for heterogeneity, as the sample variances of correlations between SNPs is high. By performing a single test over all cases and controls, we obtain a significant p-value of 8.54*e* − 4, though some heterogeneity may be contributed by batch effects. By summing scores across cohorts, we obtain a larger but still significant p-value of 0.011, suggesting that while batch effects contribute to detected heterogeneity, they do not completely account for all heterogeneity observed in the data. Lastly, by applying meta-analysis methods over individual cohort scores, we obtain a Fisher’s *χ*^2^ p-value of 0.030, and a Z-score of 2.03, also supporting the presence of a significant heterogeneity signal.

**Figure 6:**
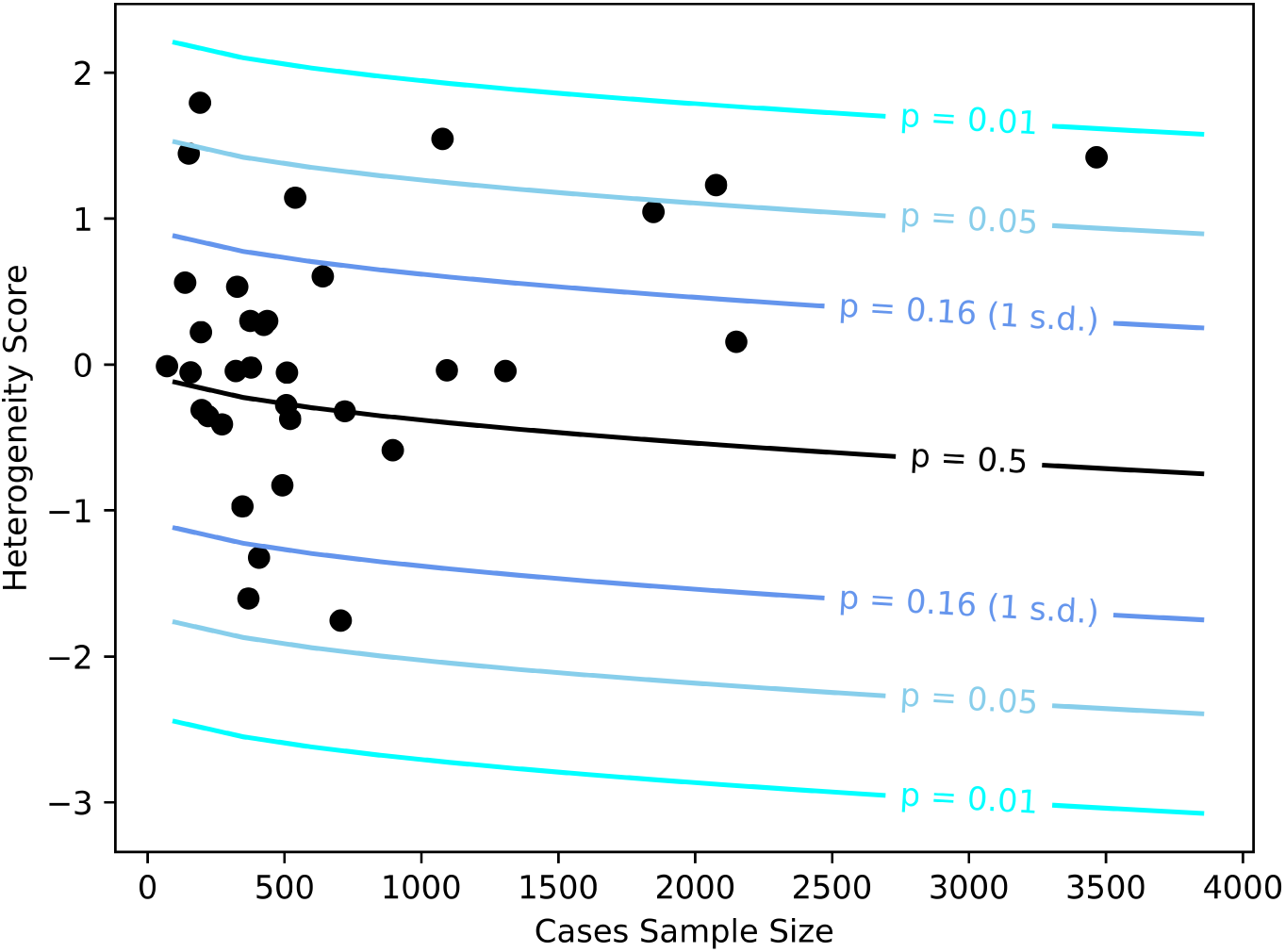
CLiP heterogeneity scores evaluated over single cohorts in the PGC schizophrenia data set, plotted by the number of genotyped cases. The black line denotes the expected score given summary statistics reported in [31] and sample sizes specific to each cohort, and the shaded region denotes z-score thresholds corresponding to particular p-values of significance.

## 4 Discussion

We present a general framework for identifying hidden heterogeneity among cases for multiple phenotypes by observing correlations between genotypes. Specifically, we derive modified test statistics to account for non-genotype input variables such as expression data, which may be continuous and sampled from any distribution with known mean and variance. Additionally, we allow for heterogeneity to be scored in quantitative phenotypes, that lack the clear-cut ascertainment of cases vs. controls that facilitates a simple dichotomous contrast of correlation patterns. Our novel, generalized framework is facilitates distinction between situations of heterogeneous subtyping and those of true pleiotropy, in which the set of individuals with each disease are themselves homogeneous.

A natural next step for these two methods is to combine them for scenarios of continuous inputs and continuous phenotypes, simultaneously. This is doable as the weighting procedures apply separately within the test statistic calculation to SNP level features and individual level features, respectively.

We further demonstrated that existing theory for heterogeneity scoring [30], assuming on a traditional multiplicative model, can be applied to more modernly accepted logistic- and liability-threshold-models. We showed that the existing score assumes independence between SNPs that is absent from modern models. It therefore fails to account for a negative bias in the score due to negatively correlated SNPs, implying an excess of false negatives and motivating recalibration.

The real data results presented in this manuscript only consider PRS based on SNPs whose association signals are genomewide significant. This removes concerns of false positive associations within the PRS. PRS constructions that do include lower-significance SNPs explain more heritability, and are an attractive next challenge for finding heterogeneity signals. Aside for the statistical challenge, this future work would require handling much larger sets of SNPs, and therefore matrices of correlations. Application of CLiP-Y to real quantitative GWAS, and CLiP-X to TWAS is still limited by related issues of small association signals or low variance explained by significant predictors. CLiP-X with measured expression data requires larger cohorts that typically assembled, as mega-analysis is often hampered by batch effects.

Given the above extensions to the correlation-based framework, it can now be applied broadly across many different traits to look for genotypic heterogeneity among cases in diseases with previously reported pleiotropic effects. While detection of heterogeneity signifies that the involved SNPs cannot be considered strictly pleiotropic, heterogeneity also suggests that distinct subgroups of significant size for a separate phenotype must exist among the cases for a particular primary phenotype under study. As incidence rates for most diseases are low, detection of significant heterogeneity may suggest a higher degree of comorbidity than is expected at random. Therefore, among disease pairs for which heterogeneity is discovered, identifying the particular subgroups underlying an elevated heterogeneity score may lead to further insights into pleiotropic interactions between phenotypes. Lastly, this framework presents the possibility of screening a large number of potentially pleiotropic secondary phenotypes against a single primary phenotype of interest. All that is required is a cohort pertaining to the primary phenotype, and a set of SNPs associated with the second phenotype whose correlations are to be evaluated over the cohort.

At the grander scheme of human genetics, generalized testing for heterogeneity paves the way for recovering additional layers of the network of effects that explain traits by interacting genetic and other factors. Going beyond the the first-order, linear approximation of these effects holds the promise of better explaining mechanisms beyond identification of their contributing input factors.

## 6 Acknowledgements

This work was supported in part by grant R01 MH117646-02S1 from the National Institutes of Health to T.L. Itsik Pe’er is supported by grants CCF-1547120 and DGE-1144854 from the National Science Foundation as well as grant U54CA209997 from the National Institutes of Health.

## 7 Web Resources

A Github page for CLiP, including code to reproduce figures is available at: https://github.com/jyuan1322/CLiP

