## Supplementary Material for "Leveraging correlations between polygenic risk score predictors to detect heterogeneity in GWAS cohorts"

### 8 Supplemental Material

#### 8.1 Negative correlations: an intuitive explanation

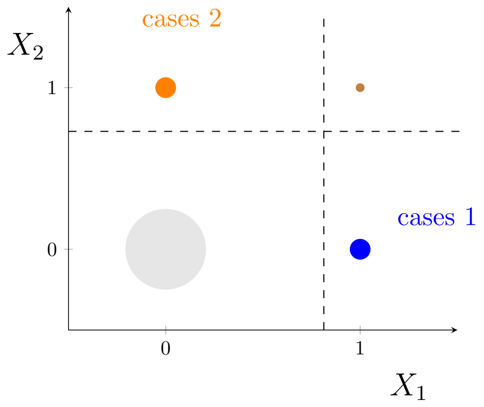


Figure S1: Correlations between PRS predictors are apparent when considering a highly simplified PRS. Assume a threshold model comprising only two independent haploid SNPs $X_1$ and $X_2$, and $\beta_1 = \beta_2 = 1$; i.e. an individual with a single risk-allele for either SNP is classified as a case. The probability of having $X_1 = 1$ given an individual is a case decreases if it is known that $X_2 = 1$. This results in a negative correlation between the two variables.

We demonstrate that among cases that are selected based on a thresholded linear score, as in logistic or liability threshold GWAS models, the correlations between SNPs is expected to be nonzero if those SNPs contribute a nonzero effect to determining the GWAS phenotype. Intuitively, this can be observed in an extreme scenario in which cases and controls are determined by two variables, $X_1$ and $X_2$, with a variance explained of 1. This scenario is visualized in S1A. We would like to evaluate the probability of an individual possessing $X_1 = 1$ given that that individual is a case. In the absence of any knowledge of $X_2$, there approximately an equal chance that the individual is a case because $X_1 = 1$ or $X_2 = 1$, with a typically negligible probability that both are 1.

$$P(\text{case } 1 | \text{case } 1 \cup \text{case } 2) =$$

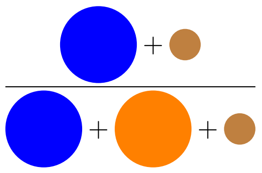

$$\approx 0.5 \qquad (24)$$

However, if it is known that a case has $X_2 = 1$, the case must inhabit the region above the threshold line on the $X_2$ axis, and so $X_1 = 1$ can only be satisfied if the individual belongs to one of the rare cases with both variables equal to 1.

$$P(\text{case } 1|\text{case } 2) =$$

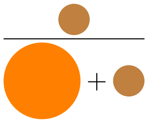

$$\ll 0.5 \tag{25}$$

Therefore in this extreme scenario $X_1$ and $X_2$ are negatively correlated. Had one of the variables decreased the risk score of the individual instead, then by the same logic $X_1$ and $X_2$ would be positively correlated. A full proof describing more realistic scenarios in which there may be a large number of small effects can be found in the Supplemental Note.

#### 8.2 Associated SNPs are correlated among cases in logistic and liability threshold models

Assume there exists a logistic model for disease risk with effects (log odds ratios) for associated SNPs $\{\beta_1, ..., \beta_M\}$. Further, assume we have collected a sample of $N$ cases of a case/control study for which the model predicts disease risk, represented by genotype matrix $X \in \{0, 1, 2\}^{N \times M}$ and labels $Y = \{1\}^N$. We demonstrate that when taken over the matrix $X$, the correlation $r$ between any two SNPs with nonzero effect is nonzero. Specifically, given two SNPs $j$ and $k$ with effects $\beta_j$ and $\beta_k$,

1. $\beta_j = 0 \cup \beta_k = 0 \implies r_{jk} = 0$
2. $\text{sign}(\beta_j) = \text{sign}(\beta_k) \implies r_{jk} < 0$
3. $\text{sign}(\beta_j) \neq \text{sign}(\beta_k) \implies r_{jk} > 0$

The correlation between SNPs $j$ and $k$ in the sample, represented as $X_j$ and $X_k$, is by definition

$$r_{jk} = \frac{\mathbb{E}[X_j X_k] - \mathbb{E}[X_j]\mathbb{E}[X_k]}{\sqrt{\mathbb{E}[X_j^2] - \mathbb{E}[X_j]^2}\sqrt{\mathbb{E}[X_k^2] - \mathbb{E}[X_k]^2}} \tag{26}$$

and so the sign of $r_{jk}$ is determined by the sign of the numerator. Here $E[X_j]$ represents the expected risk-allele count of SNP $j$ within the set of cases. Note that in a multiplicative Risch model this numerator is always 0. With a disease prevalence $B$ and odds ratios $\{b_1, ..., b_M\}$, the probability of generating a case given the number of risk-alleles for SNP $i$ is $x_i$ is $P(y = 1|x_i) = Bb_i^{x_i}D$ where $D = \prod_{l=\{1,...,M\}_{-i}} b_l^{x_l}$ is the expected contribution to the posterior of the remaining $M - 1$ SNPs. The expectation of risk-allele counts can be derived from these posteriors:

$$E[x_i] = \sum_{x\in\{0,1,2\}} x \frac{BDb_i^x p_i(x)}{\sum_{t\in\{0,1,2\}} BDb_i^t p_i(t)} = \sum_{x\in\{0,1,2\}} x \frac{a_i(x)}{\sum_{t\in\{0,1,2\}} a_i(t)} \tag{27}$$

where $a_i(x) = b_i^x p_i(x)$. If the expectation of a product of two SNPs is expressed in the same manner, it is clearly equivalent to the product of the expectation of each SNP.

$$\begin{aligned} E[x_i x_j] &= \sum_{x_i\in\{0,1,2\}} \sum_{x_j\in\{0,1,2\}} x_i x_j \frac{a_i(x_i)a_j(x_j)}{\sum_{t_i\in\{0,1,2\}} \sum_{t_j\in\{0,1,2\}} a_i(t_i)a_j(t_j)} \\ &= \frac{1}{\sum_{t_i\in\{0,1,2\}} a_i(t_i)} \sum_{x_i\in\{0,1,2\}} x_i a_i(x_i) \frac{1}{\sum_{t_j\in\{0,1,2\}} t_j a_j(t_j)} \sum_{x_j\in\{0,1,2\}} x_j a_j(x_j) \\ &= E[x_i]E[x_j] \end{aligned} \tag{28}$$

Therefore effect SNPs in the multiplicative Risch model are uncorrelated in cases.

Using the law of total expectation, rewrite the expectation of the product of $X_j$ and $X_k$ as an expectation over $X_j$ conditional on $X_k$:

$$\begin{aligned} \mathbb{E}[X_j X_k] &= \mathbb{E}_{X_k}[\mathbb{E}[X_j X_k | X_k]] \\ &= \mathbb{E}[X_k \mathbb{E}[X_j | X_k]] \end{aligned} \tag{29}$$

This term is substituted into the numerator of the correlation in equation 26 and expanded over the marginalization of $X_k = \{0, 1, 2\}$, with the $X_k = 0$ term canceling.

$$
\begin{aligned}
\mathbb{E}[X_jX_k] - \mathbb{E}[X_j]\mathbb{E}[X_k] =& \mathbb{E}[X_k\mathbb{E}[X_j|X_k]] - \mathbb{E}[X_k\mathbb{E}[X_j]] \\
=& \Bigg[p(X_k = 1)\mathbb{E}[X_j|X_k = 1] + 2p(X_k = 2)\mathbb{E}[X_j|X_k = 2]\Bigg] - \\
& \Bigg[p(X_k = 1)\mathbb{E}[X_j] + 2p(X_j = 2)\mathbb{E}[X_j]\Bigg]
\end{aligned} \tag{30}
$$

This expression determines the sign of the correlation between SNPs $X_j$ and $X_k$. The expectation $E[X_j]$ can be expressed as a sum of conditional expectations on $X_j$. After rearranging terms:

$$
\begin{aligned}
\mathbb{E}[X_jX_k] - \mathbb{E}[X_j]\mathbb{E}[X_k] =& p(X_k = 1)\Bigg[\mathbb{E}[X_j|X_k = 1] - \begin{bmatrix} \mathbb{E}[X_j|X_k = 0]p(X_k = 0)+ \\ \mathbb{E}[X_j|X_k = 1]p(X_k = 1)+ \\ \mathbb{E}[X_j|X_k = 2]p(X_k = 2) \end{bmatrix}\Bigg] + \\
& 2p(X_k = 2)\Bigg[\mathbb{E}[X_j|X_k = 2] - \begin{bmatrix} \mathbb{E}[X_j|X_k = 0]p(X_k = 0)+ \\ \mathbb{E}[X_j|X_k = 1]p(X_k = 1)+ \\ \mathbb{E}[X_j|X_k = 2]p(X_k = 2) \end{bmatrix}\Bigg] \\
=& p(X_k = 1)\Bigg[\mathbb{E}[X_j|X_k = 1](1 - p(X_k = 1)) - \begin{bmatrix} \mathbb{E}[X_j|X_k = 0]p(X_k = 0)+ \\ \mathbb{E}[X_j|X_k = 2]p(X_k = 2) \end{bmatrix}\Bigg] + \\
& 2p(X_k = 2)\Bigg[\mathbb{E}[X_j|X_k = 2](1 - p(X_k = 2)) - \begin{bmatrix} \mathbb{E}[X_j|X_k = 0]p(X_k = 0)+ \\ \mathbb{E}[X_j|X_k = 1]p(X_k = 1) \end{bmatrix}\Bigg] \\
=& p(X_k = 1)\Bigg[\mathbb{E}[X_j|X_k = 1]\left(\begin{matrix} p(X_k = 0)+ \\ p(X_k = 2 \end{matrix}\right)) - \begin{bmatrix} \mathbb{E}[X_j|X_k = 0]p(X_k = 0)+ \\ \mathbb{E}[X_j|X_k = 2]p(X_k = 2) \end{bmatrix}\Bigg] + \\
& 2p(X_k = 2)\Bigg[\mathbb{E}[X_j|X_k = 2]\left(\begin{matrix} p(X_k = 0)+ \\ p(X_k = 1) \end{matrix}\right) - \begin{bmatrix} \mathbb{E}[X_j|X_k = 0]p(X_k = 0)+ \\ \mathbb{E}[X_j|X_k = 1]p(X_k = 1) \end{bmatrix}\Bigg]
\end{aligned} \tag{31}
$$

After collecting the conditional expectation terms:

$$\begin{aligned}\mathbb{E}[X_jX_k]-\mathbb{E}[X_j]\mathbb{E}[X_k]=&\mathbb{E}[X_j|X_k=0]\big(-p(X_k=0)p(X_k=1)-2p(X_k=2)p(X_k=0)\big)+\\&\mathbb{E}[X_j|X_k=1]\begin{pmatrix}p(X_k=1)p(X_k=0)+p(X_k=1)p(X_k=2)-\\2p(X_k=2)p(X_k=1)\end{pmatrix}+\\&\mathbb{E}[X_j|X_k=2]\begin{pmatrix}-p(X_k=1)p(X_k=2)+2p(X_k=2)p(X_k=0)+\\2p(X_k=2)p(X_k=1)\end{pmatrix}\end{aligned}\tag{32}$$

Probabilities of $X_k$ can be expressed as a function of the risk-allele frequency $p_k$.

$$p(X_k=0)=(1-p_k)^2 \qquad p(X_k=1)=2p_k(1-p_k) \qquad p(X_k=2)=p_k^2 \tag{33}$$

After substitution of these expressions into equation 32, the result is a sum of polynomial functions of $p_k$.

$$\begin{aligned}\mathbb{E}[X_jX_k]-\mathbb{E}[X_j]\mathbb{E}[X_k]=&\mathbb{E}[X_j|X_k=0]\big(-2p_k^3+4p_k^2-2p_k\big)+\\&\mathbb{E}[X_j|X_k=1]\big(4p_k^3-6p_k^2+2p_k\big)+\\&\mathbb{E}[X_j|X_k=2]\big(-2p_k^3+2p_k^2\big)\end{aligned}\tag{34}$$

Note that when $\mathbb{E}[X_j|X_k=0]=\mathbb{E}[X_j|X_k=0]=\mathbb{E}[X_j|X_k=0]$, equation 34 reduces to 0 for all values of $p_k$. When $\mathbb{E}[X_j|X_k=0]<\mathbb{E}[X_j|X_k=0]<\mathbb{E}[X_j|X_k=0]$, equation 34 is strictly positive over the $[0,1]$ domain of $p_k$ whereas when $\mathbb{E}[X_j|X_k=0]>\mathbb{E}[X_j|X_k=0]>\mathbb{E}[X_j|X_k=0]$, the function is strictly negative over the domain. Next we must prove that the former inequality would result if the signs of the effect sizes of $X_j$ and $X_k$ were different, whereas the latter inequality would result if the signs were the same.

For any value of $X_k$, the conditional expectation of $X_j$ can be expanded:

$$\mathbb{E}[X_j|X_k] = \frac{\sum_{X_j \in \{0,1,2\}} X_j p(Y=1|X_j, X_k)p(X_j)}{\sum_{X_j \in \{0,1,2\}} p(Y=1|X_j, X_k)p(X_j)}$$

$$= \frac{p(Y=1|X_j=1, X_k)p(X_j=1) + 2p(Y=1|X_j=2, X_k)p(X_j=2)}{p(Y=1|X_j=0, X_k)p(X_j=0) + p(Y=1|X_j=1, X_k)p(X_j=1) + p(Y=1|X_j=2, X_k)p(X_j=2))} \tag{35}$$

The probability $p(Y=1|X_j, X_k)$ is the probability of an individual being a case and is determined by a logistic function with fixed values for $X_j$ and $X_k$. The numerator of Equation 35 differs from the denominator by replacing $p(Y=1|X_j=0, X_k)p(X_j=0)$ with a second $p(Y=1|X_j=2, X_k)p(X_j=2)$ term. Consider the expression $\frac{p(Y=1|X_j=2,X_k)p(X_j=2)}{p(Y=1|X_j=0,X_k)p(X_j=0)}$. We would like to show first that the magnitude of this term decreases as $X_k \in \{0,1,2\}$ increases. Express the probabilities as logistic functions

$$\frac{\frac{1}{1+\exp\left(-(2b_j+X_k b_k+\mathbb{E}[Xb]_{-jk})\right)}}{\frac{1}{1+\exp\left(-(X_k b_k+\mathbb{E}[Xb]_{-jk})\right)}} \frac{p(X_j=2)}{p(X_k=0)} = \frac{1+\exp\left(-(X_k b_k+\mathbb{E}[Xb]_{-jk})\right)}{1+\exp\left(-(2b_j+X_k b_k+\mathbb{E}[Xb]_{-jk})\right)} \frac{p(X_j=2)}{p(X_k=0)} \tag{36}$$

where $b_j$ and $b_k$ are the odds ratios of SNPs $j$ and $k$, respectively, and $\mathbb{E}[Xb]_{-jk}$ is the expected contribution of the remaining undetermined SNPs when SNPs $j$ and $k$ are fixed. If the effect size of $X_k$ is positive, the contribution of a particular value of $X_k$ can be represented as adding a constant term $c$ within a separate exponential term.

$$
\begin{aligned}
\frac{1+\exp(-(\mathbb{E}[Xb]_{-jk}+c))}{1+\exp(-(2b_j+\mathbb{E}[Xb]_{-jk}+c))}\frac{p(X_j=2)}{p(X_k=0)} =& \frac{1+\exp(-(a+c))}{1+\exp(-(b+a+c))}D \\
=& \frac{1+\exp(-a)\exp(-c)}{1+\exp(-(b+a))\exp(-c)}D \\
=& \frac{\big[1+\exp(-a)\big]\exp(-c)+1-\exp(-c)}{\big[1+\exp(-(b+a))\big]\exp(-c)+1-\exp(-c)}D \\
=& \frac{\big[1+\exp(-a)\big]+\exp(c)-1}{\big[1+\exp(-(b+a))\big]+\exp(c)-1}D \\
=& \frac{\big(1+\exp(-\mathbb{E}[Xb]_{-jk})\big)+\exp(c)-1}{\big(1+\exp(-(2b_j+\mathbb{E}[Xb]_{-jk}))\big)+\exp(c)-1}\frac{p(X_j=2)}{p(X_k=0)}
\end{aligned}
\tag{37}
$$

Therefore, fixing $X_k$ to a particular value has the effect of adding a $\exp(c)-1$ term to the numerator and denominator of the ratio. Additionally, as the number of risk-alleles of $X_k$ is increased, the value of $c$ and thus $\exp(c)-1$ increases. This allows us to take advantage of the following Lemma:

**Lemma:** for strictly positive values of $c, d, e$ with $c > d$, $\frac{c}{d} > \frac{c+e}{d+e}$. $c > d$ is satisfied for all conditional expectations of $X_j$ provided that the effect of $X_j$ is positive.

$$
\begin{aligned}
c >& d \\
ce + cd >& de + cd \\
c(e+d) >& d(e+c) \\
\frac{c}{d} >& \frac{c+e}{d+e}
\end{aligned}
\tag{38}
$$

Now substitute the fraction $\frac{p(Y=1|X_j=2,X_k)p(X_j=2)}{p(Y=1|X_j=0,X_k)p(X_j=0)}$ for $\frac{c}{d}$. When the value of $X_k$ is fixed, the change to the probability is equivalent to adding a constant $e$ to both the numerator and denominator, as shown in Equation 37. By the above lemma, an increase in $X_k$ and thus an increase in the magnitude of $e$ reduces the value of $\frac{p(Y=1|X_j=2,X_k)p(X_j=2)}{p(Y=1|X_j=0,X_k)p(X_j=0)}$ as $\frac{c}{d}$.

Next, rewrite the fraction as $p(Y=1|X_j=2,X_k)p(X_j=2) = ap(Y=1|X_j=0,X_k)p(X_j=0)$

where $a$ is some positive constant greater than 1, as with all other variables fixed and a positive effect $b_j$, $X_j = 2$ will always increase the probability of generating a case vs $X_j = 0$. We can then eliminate $p(Y = 1|X_j = 2, X_k)p(X_j = 2)$ in Equation 35. For simplicity we have also made the substitution $p_x = p(Y = 1|X_j = x, X_k)p(X_j = x)$:

$$\begin{aligned}
\mathbb{E}[X_j|X_k] =& \frac{\begin{array}{c} p(Y=1|X_j=1,X_k)p(X_j=1)+ \\ 2p(Y=1|X_j=2,X_k)p(X_j=2)\end{array}}{\begin{array}{c} p(Y=1|X_j=0,X_k)p(X_j=0)+ \\ p(Y=1|X_j=1,X_k)p(X_j=1)+ \\ p(Y=1|X_j=2,X_k)p(X_j=2))\end{array}} \\
=& \frac{p_1 + 2p_2}{p_0 + p_1 + p_2} \\
=& \frac{p_1 + 2ap_0}{p_1 + (1+a)p_0}
\end{aligned} \tag{39}$$

The first and second derivatives of this function with respect to $a$ are:

$$\begin{aligned}
\frac{d}{da}\left[\frac{p_1 + 2ap_0}{p_1 + (1+a)p_0}\right] =& \frac{p_0(2p_0 + p_1)}{(p_0 + p_1 + p_0 a)^2} \\
\frac{d^2}{da^2}\left[\frac{p_1 + 2ap_0}{p_1 + (1+a)p_0}\right] =& \frac{-2p_0^2(2p_0 + p_1)}{(p_0 + p_1 + p_0 a)^3}
\end{aligned} \tag{40}$$

For positive values of $a$, the first derivative is always positive, and the second derivative is always negative, indicating that this is a monotonic, concave function. We have previously established that as $X_k$ increases in magnitude, $a$ decreases; that is, $a_{k=0} > a_{k=1} > a_{k=2}$. As $E[X_j|X_k]$ is a monotonically increasing function of $a$, then the previous inequality is likewide true for $E[X_j|X_k]$. Therefore, $\mathbb{E}[X_j|X_k = 0, Y = 1] > \mathbb{E}[X_j|X_k = 1, Y = 1] > \mathbb{E}[X_j|X_k = 2, Y = 1]$. Conversely, when $X_j$ has an effect size less than 0, and thus opposite that of $X_j$, then $\mathbb{E}[X_j|X_k = 0, Y = 1] < \mathbb{E}[X_j|X_k = 1, Y = 1] < \mathbb{E}[X_j|X_k = 2, Y = 1]$.

#### 8.3 Derivation of heterogeneity score with continuous inputs

Starting with the expression for correlation weighted by $\pi$,

$$r_{jk} = \frac{\begin{array}{c}\pi\mathbb{E}\big[X_j^+ X_k^+\big] + (1-\pi)\mathbb{E}\big[X_j^- X_k^-\big] - \\ (\pi\mu_j^+ + (1-\pi)\mu_j^-)(\pi\mu_k^+ + (1-\pi)\mu_k^-)\end{array}}{\sqrt{\begin{array}{l}\pi\mathbb{E}\big[(X_j^+)^2\big] + \\ \quad (1-\pi)\mathbb{E}\big[(X_j^-)^2\big] - \\ \qquad \big[\pi\mu_j^+ + (1-\pi)\mu_j^-\big]^2\end{array}}\sqrt{\begin{array}{l}\pi\mathbb{E}\big[(X_k^+)^2\big] + \\ \quad (1-\pi)\mathbb{E}\big[(X_k^-)^2\big] - \\ \qquad \big[\pi\mu_k^+ + (1-\pi)\mu_k^-\big]^2\end{array}}} \tag{41}$$

Taking the derivative with respect to $\pi$ at $\pi = 0$ allows the cancellation of many terms. If $r_{jk}$ is expressed as $r_{jk} = \frac{a}{bc}$, where $a$ is the entire numerator and $b$ and $c$ are the two standard deviation terms in the denominator, then $\frac{\partial}{\partial\pi} r_{jk}\big|_{\pi=0} = \frac{a'bc - a(b'c+bc')}{b^2c^2}$. With $\pi = 0$, the second term in numerator simplifies to $\mathbb{E}[X_j^- X_k^-] - \mu_j^- \mu_k^-$, which equals zero when we assume there is no correlation between SNPs in a homogeneous case or control set. Therefore we evaluate $\frac{\partial}{\partial\pi} r_{jk}\big|_{\pi=0} = \frac{a'}{bc}$.

From our assumption that there is no correlation within cases or controls

$$\mathbb{E}[X_j^+ X_k^+] = \mu_j^+ \mu_k^+ \tag{42}$$

$$\mathbb{E}[(X_j^+)^2] = (\mu_j^+)^2 + (\sigma_j^+)^2 \tag{43}$$

where $(\sigma_j^+)^2$ is the variance of SNP $j$ within sub-cohort $+$. Lastly, at $\pi = 0$, each standard deviation term in the denominator reduces to

$$\mathbb{E}[(X_j^-)^2] - (\mu_j^-)^2 = \big[(\mu_j^-)^2 + (\sigma_j^-)^2\big] - (\mu_j^-)^2 = (\sigma_j^-)^2 \tag{44}$$

This yields the heterogeneity test statistic for continuous inputs.

$$w_{jk} = \frac{\partial}{\partial \pi} r_{jk}\Big|_{\pi=0} \tag{45}$$
$$= \frac{\mu_j^+ \mu_k^+ - \mu_j^+ \mu_k^- - \mu_j^- \mu_k^+ + \mu_j^- \mu_k^-}{\sigma_j^- \sigma_k^-} \tag{46}$$
$$= \frac{(\mu_j^+ - \mu_j^-)(\mu_k^+ - \mu_k^-)}{\sigma_j^- \sigma_k^-} \tag{47}$$

#### 8.4 Prediction of heterogeneity scores in homogeneous (null) cohorts

The heterogeneity score relies on the weighted difference in correlations between cases and controls. The test for heterogeneity assumes in the null situation that a cohort of cases is completely homogeneous, i.e. sampled and thresholded using the same polygenic risk score model. Therefore expected sample correlations between every pair of predictors $X_i$ and $X_j$ (either SNP allele counts or gene expression measurements) are computed assuming that all individuals are identically sampled cases.

$$r(X_i, X_j) = \frac{\mathbb{E}[X_i X_j] - \mathbb{E}[X_i]\mathbb{E}[X_j]}{\sqrt{\mathbb{E}[X_i^2] - \mathbb{E}[X_i]^2}\sqrt{\mathbb{E}[X_j^2] - \mathbb{E}[X_j]^2}} \tag{48}$$

By Bayes theorem, each of these expectations can be computed from the posterior probabilities of the predictor values given an individual is a case:

$$\begin{aligned}
\mathbb{E}[x_i|\text{case}] =& \frac{\int x_i P(\text{case}|x_i) P(x_i) dx_i}{\int P(\text{case}|x_i = z) P(x_i = z) dz} \\
\mathbb{E}[x_i^2|\text{case}] =& \frac{\int x_i^2 P(\text{case}|x_i) P(x_i) dx_i}{\int P(\text{case}|x_i = z) P(x_i = z) dz} \\
\mathbb{E}[x_i x_j|\text{case}] =& \frac{\int \int x_i x_j P(\text{case}|x_i, x_j) P(x_i, x_j) dx_i dx_j}{\int \int P(\text{case}|x_i = z, x_j = w) P(x_i = z, x_j = w) dz dw}
\end{aligned} \tag{49}$$

The prior probability for any predictor $P(X_i)$ when $X$ are SNPs is simply the binomial distribution parameterized by the risk-allele frequency in controls. As we assume that SNPs are sampled from independent loci, their joint distributions are simply the product of these priors.

$$P(X_i) = f_c^{X_i}(1 - f_c)^{(2-X_i)} \tag{50}$$

When the $M$ predictors $X_i$ are gene expression variables, additional correlations may result if SNPs are associated with more than one gene. The prior joint distribution $P(X_i, X_j)$ in controls can be calculated given the linearity of the covariance with respect to each of the variables being correlated. First, assume a linear model over genotypes generates each quantitative transcript, and a linear model over these transcripts in turn generates the hidden liability score.

$$X_i = \beta_i G \quad \forall i \in 1, ..., M \tag{51}$$

$$y^* = \sum_{i=1}^{M} \alpha_i X_i \tag{52}$$

$$\text{Cov}(X_i, X_j) = \text{Cov}(\beta_i G, \beta_j G) = 2\alpha_i \alpha_j \sum_{s=1}^{S} \beta_{si} \beta_{sj} \text{Var}(G_s) \tag{53}$$

$$\text{Var}(G_s) = 2 f_{sc}(1 - f_{sc}) \tag{54}$$

Lastly, the probability of an individual being a case given a particular predictor value, $P(case|X_i)$, is determined by the liability threshold equation, where $\Phi$ is the normal CDF. The mean of the liability distribution is shifted by the value, and the variance explained by the predictor is subtracted from the total liability variance (standardized to 1).

$$P(y = 1|X_i = b) = 1 - \Phi\left(\frac{X_i \alpha_i - T}{\sqrt{1 - \text{Var}(X_i)}}\right) \tag{55}$$

The expectation of predictor products in the numerator of the correlation is determined by the probability of an individual being a case given a pair of predictor values. First, assume the pair of known predictors comprises the variable set $b$, and the remaining unknown predictors set $a$. The distribution over all predictors is the block multivariate normal distribution parametrized by $\mu = \left(\begin{smallmatrix} \mu_a \\ \mu_b \end{smallmatrix}\right)$ and $\Sigma = \left(\begin{smallmatrix} \Sigma_{aa} & \Sigma_{ab} \\ \Sigma_{ba} & \Sigma_{bb} \end{smallmatrix}\right)$. Given the values of $b$, the conditional distribution of the remaining predictors $a$ has the following mean and covariance [38]

$$\mu_{a|b} = \mu_a + \Sigma_{ab}\Sigma_{bb}^{-1}(X_b - \mu_b) \tag{56}$$

$$\Sigma_{a|b} = \Sigma_{aa} - \Sigma_{ab}\Sigma_{bb}^{-1}\Sigma_{ba} \tag{57}$$

The mean and variance of the liability score can be calculated as the weighted sum of these predictors, and these estimates can be substituted into the liability threshold equation as before to calculate $P(y = 1|X_i = b_1, X_j = b_2)$.

#### 8.5 Algorithms for CLiP-X and CLiP-Y Simulation and CLiP-Y polynomial weight search

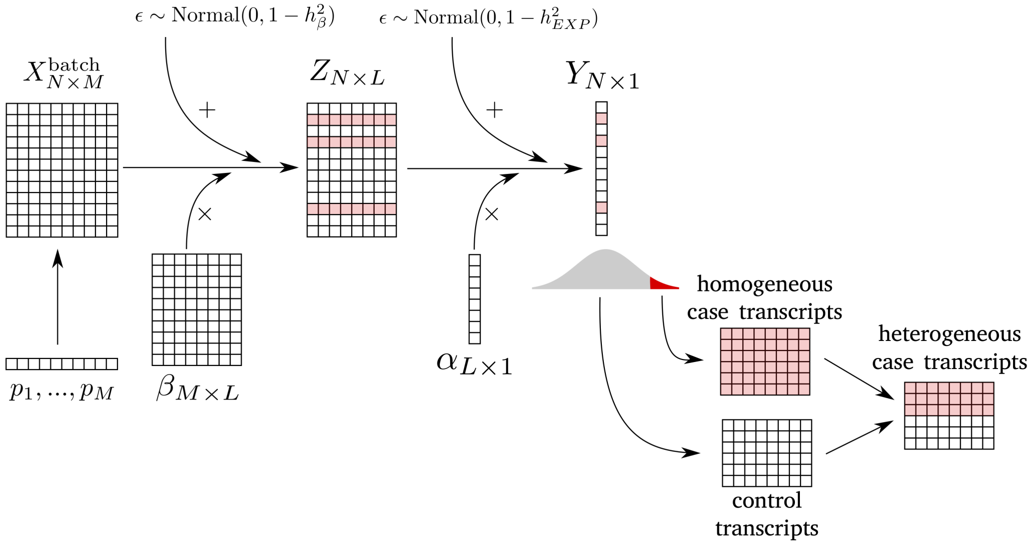


Figure S2: Sampling procedure for CLiP-X: heterogeneity detection with quantitative input variables. Because in most scenarios the inputs will represent gene expression, we simulated transcripts as linear functions of randomly sampled genotypes, allowing for prior correlations from single genotypes associated with multiple transcripts. To generate a desired number of cases (Red), batches of random transcripts are generated repeatedly, and those individuals whose liability scores pass the threshold $T$ are concatenated to a growing list of cases. Heterogeneous cases are created by concatenating true cases with controls.

**Function** SampleCLiP-X

**Input:** $N, M, L, h^2_{SNP}, h^2_{EXP}, \pi$ **Output:** genotypes $X_{N\times M}$, expression $Z_{N\times L}$

$T = \Phi^{-1}(1 - .01)$; `// threshold from 0.01 prevalence`

`/* define SNP and expression summary statistics */`

$p_{i\in[1,M]} = 0.5; \quad \beta_{i\in[1,M]} = \sqrt{\frac{h^2_{SNP}/M}{\mathrm{Var}(X_{\cdot i})}}; \quad \alpha_{i\in[1,L]} = \sqrt{\frac{h^2_{EXP}/L}{\mathrm{Var}(Z_{\cdot i})+2\sum_{j\in L}\mathrm{Cov}(Z_{\cdot i}, Z_{\cdot j})}};$

`/* Generate control genotypes and expression */`

**for** $n$ *in* $[1, (1-\pi)N]$, $m$ *in* $[1, M]$ **do**

  $X^0_{n,m} \sim \mathrm{Binom}(2, p_m)$;

**end**

$Z^0 \sim \mathrm{Normal}(X^0 \cdot \beta, 1 - h^2_{SNP})$;

`/* Generate case genotypes and expression */`

$X = []$; $Z = []$;

**while** $\mathrm{nrows}(X) = \mathrm{nrows}(Z) < \pi N$ **do**

  **for** $m$ *in* $[1, M]$ **do**

    $x_m \sim \mathrm{Binom}(2, p_m)$;

  **end**

  $z \sim \mathrm{Normal}(x \cdot \beta, 1 - h^2_{SNP})$;

  $y \sim \mathrm{Normal}(z \cdot \alpha, 1 - h^2_{EXP})$;

  **if** $y \geq T$ **then**

    $\mathrm{append}(X, x)$;

    $\mathrm{append}(Z, z)$;

  **end**

**end**

`/* Simulate heterogeneity by concatenating generated cases and controls */`

$X = \mathrm{concatenate}(X, X^0)$;

$Z = \mathrm{concatenate}(Z, Z^0)$;

**Algorithm 1:** Sampling procedure for heterogeneous cases with quantitative predictors.

**Function** SampleCLiP-Y

**Input:** $N, M, h^2_{SNP}, cohort = \{hom, het\}, \pi$

**Output:** $X_{N \times M}, Y_{N \times 1}$

`/* define SNP and expression summary statistics */`

$p_{i \in [1,M]} = 0.5; \quad \beta_{i \in [1,M]} = \sqrt{\frac{h^2_{SNP}/M}{\mathrm{Var}(X_{\cdot i})}};$

**for** $n$ *in* $[1, N]$, $m$ *in* $M$ **do**

  $X_{nm} \sim \mathrm{Binom}(2, p_m);$

**end**

`/* if heterogeneous, generate mislabeled individuals */`

**if** $cohort = het$ **then**

  $Y_{[1,N/2]} \sim \mathrm{Normal}(X_{[1,N/2],\cdot} \cdot \beta, 1 - h^2_{SNP});$

  $Y^0_{[1,N/2]} \sim \mathrm{Normal}(\mathbb{E}[X\beta], 1 - h^2_{SNP});$

  $Y = \mathrm{concatenate}(Y, Y^0)$

**else**

  $Y \sim \mathrm{Normal}(X \cdot \beta, 1 - h^2_{SNP});$

**end**

**Algorithm 2:** Sampling procedure for heterogeneous PRS cohorts with quantitative phenotypes. In lieu of explicit case/control labels by which to generate heterogeneous cohorts, we define 'cases' as individuals whose phenotypes are a function of their PRSs, whereas 'controls' have phenotypes that are sampled completely randomly from the same distribution.

**Function** PolynomialWeightSearch

**Input:**

$deg$; `// Polynomial degree`

$N$; `// Size of the case/control cohort to simulate`

$k_{disc} = 5$, $k_{val} = 20$; `// Number sample cohorts to simulate for discovery, validation steps`

$S_{thresh}$; `// Threshold score for saving candidate coefficient`

coef_sd; `// std. dev. for random increments in coefficient search`

**Output:** $coefs$

**Function** `CalScore`($\{X_{het}\}, \{X_{hom}\}, coefs$):

  **return** Mean(CLiP-Y($\{X_{het}\}, coefs$)) − Mean(CLiP-Y($\{X_{hom}\}, coefs$));

**Function** `DiscoverCandidatePolynomials`($\{X_{het}\}, \{X_{hom}\}, S_{thresh}$):

  $cands = \{\}$ **for** Num candidates desired **do**

    $coefs = \mathbb{R}_{(deg+1)} \sim N(0, \text{coef\_sd})$;

    $S_{het} =$ **CalcScore**($\{X_{het}\}, \{X_{hom}\}, coefs$);

    **while** Not converged **do**

      increment a randomly selected coefficient in $coefs$ by $a \sim N(0, \text{coef\_sd})$;

      **if** ***CalcScore***($\{X_{het}\}, \{X_{hom}\}, coefs$) $> S_{het}$ **then**

        update $coefs$ and $S_{het}$

      **end**

    **end**

    **if** $S_{het} > S_{thresh}$ **then**

      store $coefs$ in $cands$

    **end**

  **end**

  return $cands$

`/* Simulate cohorts for discovery/validation */`

$\{X_{het}\}_{disc} = \{X_1, ..., X_{k_{disc}} | X_i = \text{SampleCLiP-Y}(N, cohort = het)\}$;

$\{X_{het}\}_{disc} = \{X_1, ..., X_{k_{disc}} | X_i = \text{SampleCLiP-Y}(N, cohort = hom)\}$;

$cands =$ **DiscoverCandidatePolynomials**($\{X_{het}\}_{disc}, \{X_{hom}\}_{disc}, S_{thresh}$);

$\{X_{het}\}_{val} = \{X_1, ..., X_{k_{val}} | X_i = \text{SampleCLiP-Y}(N, cohort = het)\}$;

$\{X_{het}\}_{val} = \{X_1, ..., X_{k_{val}} | X_i = \text{SampleCLiP-Y}(N, cohort = hom)\}$;

**return** $\text{argmax}_{coefs \in cands}$(**CalcScore**($\{X_{het}\}_{val}, \{X_{hom}\}_{val}, coefs$))

**Algorithm 3:** Local search for optimal polynomial weight function $\phi$ for use with CLiP-Y. This weight function is applied over individuals according to their phenotypes as a percentile of the phenotype distribution in the domain $[0, 1]$, and determines the contribution of individuals to the weighted correlation substitution in CLiP-Y. Optimization is performed over a small set of discovery simulated cohorts, and validated in a larger set of simulated cohorts. A set of candidate functions which pass a fixed threshold are stored, and of these the function which performs best on the larger validation set is selected.

#### 8.6 Additional simulation results for misclassified and disjoint heterogeneity

| % SNPs shared / # sub-phenos | 0 | 0.25 | 0.5 | 0.75 | 1 |
|---|---|---|---|---|---|
| 1 | $-5.08 \pm 1.01$ | $-4.92 \pm 1.07$ | $-4.93 \pm 0.85$ | $-5.16 \pm 0.96$ | $-5.32 \pm 0.91$ |
| 2 | $-1.67 \pm 1.12$ | $-1.87 \pm 1.07$ | $-2.60 \pm 0.67$ | $-3.67 \pm 0.91$ | $-5.41 \pm 1.12$ |
| 3 | $-0.65 \pm 0.79$ | $-1.23 \pm 0.91$ | $-1.98 \pm 0.90$ | $-2.90 \pm 0.74$ | $-4.99 \pm 0.78$ |
| 4 | $-0.49 \pm 0.95$ | $-0.61 \pm 1.07$ | $-1.33 \pm 1.01$ | $-2.74 \pm 0.88$ | $-5.48 \pm 0.95$ |
| 5 | $-0.56 \pm 1.00$ | $-0.81 \pm 1.01$ | $-0.97 \pm 1.02$ | $-2.00 \pm 1.23$ | $-5.53 \pm 0.80$ |
| 6 | $-0.44 \pm 1.09$ | $-0.39 \pm 1.01$ | $-1.14 \pm 0.93$ | $-2.05 \pm 0.81$ | $-5.07 \pm 0.75$ |
| 7 | $-0.05 \pm 0.90$ | $-0.12 \pm 0.98$ | $-0.43 \pm 0.92$ | $-1.68 \pm 0.93$ | $-5.42 \pm 1.04$ |
| 8 | $0.37 \pm 0.81$ | $-0.27 \pm 0.61$ | $-0.40 \pm 1.07$ | $-1.33 \pm 0.91$ | $-5.08 \pm 1.07$ |

Table S1: Simulated CLiP results using cases generated from multiple correlated sub-phenotypes. Entries comprise mean and standard deviations of CLiP scores evaluated over 20 trials, with a total variance explained of 0.05 and total case cohort size of 50000. All simulations were performed with 100 SNPs, and in independent sub-phenotypes these 100 were subdivided equally among the number of sub-phenotypes. The percentage of SNPs shared refers to the percentage of SNPs within each sub-phenotype which has a fixed effect size across all sub-phenotypes.

| # cases \| % true ($h^2$) | | 0.01 | 0.025 | 0.05 | 0.075 | 0.1 |
|---|---|---|---|---|---|---|
| 5000 | 0 | $0.21 \pm 1.08$ | $0.07 \pm 0.85$ | $-0.16 \pm 0.72$ | $-0.21 \pm 0.91$ | $-0.07 \pm 0.79$ |
| | 0.25 | $0.14 \pm 0.64$ | $0.18 \pm 0.87$ | $1.13 \pm 0.85$ | $1.30 \pm 0.97$ | $2.00 \pm 0.80$ |
| | 0.5 | $-0.02 \pm 0.94$ | $0.80 \pm 0.98$ | $1.50 \pm 0.88$ | $2.58 \pm 1.11$ | $3.87 \pm 0.93$ |
| | 0.75 | $0.11 \pm 1.01$ | $0.53 \pm 0.90$ | $0.73 \pm 1.00$ | $1.24 \pm 1.17$ | $1.49 \pm 0.90$ |
| | 1.0 | $-0.25 \pm 0.88$ | $-0.94 \pm 0.94$ | $-1.63 \pm 0.78$ | $-2.42 \pm 0.91$ | $-3.49 \pm 1.05$ |
| 10000 | 0 | $-0.08 \pm 0.90$ | $-0.20 \pm 1.01$ | $0.21 \pm 0.97$ | $0.41 \pm 0.99$ | $-0.11 \pm 1.16$ |
| | 0.25 | $-0.34 \pm 0.62$ | $0.63 \pm 0.74$ | $1.10 \pm 1.03$ | $3.01 \pm 0.88$ | $4.07 \pm 0.84$ |
| | 0.5 | $0.28 \pm 0.83$ | $1.07 \pm 1.32$ | $2.23 \pm 1.04$ | $4.17 \pm 0.63$ | $6.13 \pm 0.99$ |
| | 0.75 | $-0.02 \pm 1.19$ | $0.95 \pm 1.23$ | $1.75 \pm 0.69$ | $2.04 \pm 0.88$ | $2.63 \pm 0.99$ |
| | 1.0 | $-0.31 \pm 0.74$ | $-0.75 \pm 1.11$ | $-2.38 \pm 1.05$ | $-3.52 \pm 1.04$ | $-4.79 \pm 1.04$ |
| 25000 | 0 | $0.19 \pm 1.08$ | $-0.25 \pm 0.99$ | $0.30 \pm 1.14$ | $0.28 \pm 0.94$ | $0.12 \pm 1.05$ |
| | 0.25 | $0.82 \pm 0.82$ | $1.47 \pm 0.74$ | $3.27 \pm 0.91$ | $5.55 \pm 1.15$ | $7.33 \pm 1.33$ |
| | 0.5 | $0.43 \pm 1.11$ | $2.06 \pm 0.98$ | $5.15 \pm 1.10$ | $7.27 \pm 0.92$ | $9.73 \pm 1.08$ |
| | 0.75 | $0.59 \pm 1.00$ | $1.05 \pm 0.97$ | $2.28 \pm 1.13$ | $3.32 \pm 0.94$ | $4.72 \pm 1.05$ |
| | 1.0 | $-0.75 \pm 0.84$ | $-1.94 \pm 1.19$ | $-3.55 \pm 0.89$ | $-5.45 \pm 1.00$ | $-7.82 \pm 0.83$ |
| 50000 | 0 | $-0.23 \pm 1.07$ | $-0.12 \pm 0.71$ | $0.21 \pm 0.81$ | $0.43 \pm 1.15$ | $-0.15 \pm 1.04$ |
| | 0.25 | $0.56 \pm 1.08$ | $2.22 \pm 0.78$ | $5.29 \pm 1.11$ | $8.89 \pm 1.36$ | $11.78 \pm 1.07$ |
| | 0.5 | $1.15 \pm 1.17$ | $3.35 \pm 0.93$ | $7.01 \pm 1.00$ | $10.69 \pm 1.28$ | $13.86 \pm 0.94$ |
| | 0.75 | $0.95 \pm 0.99$ | $1.89 \pm 0.83$ | $3.66 \pm 0.75$ | $5.19 \pm 0.76$ | $6.40 \pm 1.02$ |
| | 1.0 | $-0.93 \pm 1.06$ | $-2.51 \pm 1.10$ | $-5.10 \pm 0.82$ | $-8.43 \pm 0.99$ | $-10.74 \pm 0.96$ |
| 100000 | 0 | $0.39 \pm 0.94$ | $-0.43 \pm 1.02$ | $-0.05 \pm 0.80$ | $0.15 \pm 0.95$ | $0.15 \pm 0.95$ |
| | 0.25 | $1.17 \pm 0.99$ | $3.50 \pm 1.20$ | $8.59 \pm 1.16$ | $12.78 \pm 1.23$ | $17.05 \pm 0.84$ |
| | 0.5 | $1.61 \pm 1.09$ | $5.20 \pm 0.95$ | $10.07 \pm 1.18$ | $15.31 \pm 1.11$ | $20.76 \pm 1.07$ |
| | 0.75 | $0.66 \pm 0.74$ | $2.42 \pm 0.81$ | $4.56 \pm 1.01$ | $7.10 \pm 1.11$ | $9.04 \pm 1.08$ |
| | 1.0 | $-1.48 \pm 1.09$ | $-3.68 \pm 1.14$ | $-7.44 \pm 0.97$ | $-11.21 \pm 1.00$ | $-15.05 \pm 0.78$ |
| 250000 | 0 | $-0.18 \pm 0.99$ | $0.15 \pm 0.99$ | $-0.04 \pm 0.71$ | $0.37 \pm 0.89$ | $-0.40 \pm 1.04$ |
| | 0.25 | $2.66 \pm 0.83$ | $6.36 \pm 1.19$ | $13.51 \pm 0.99$ | $21.10 \pm 0.78$ | $27.63 \pm 0.85$ |
| | 0.5 | $3.35 \pm 1.13$ | $8.29 \pm 0.83$ | $15.99 \pm 1.04$ | $24.57 \pm 1.21$ | $32.30 \pm 1.03$ |
| | 0.75 | $1.43 \pm 0.82$ | $3.79 \pm 0.73$ | $7.61 \pm 0.83$ | $10.94 \pm 1.09$ | $15.00 \pm 1.15$ |
| | 1.0 | $-2.10 \pm 1.11$ | $-5.79 \pm 1.21$ | $-11.56 \pm 0.75$ | $-18.16 \pm 0.95$ | $-24.52 \pm 0.84$ |
| 500000 | 0 | $0.51 \pm 0.92$ | $0.23 \pm 0.70$ | $-0.06 \pm 0.97$ | $-0.21 \pm 0.75$ | $0.04 \pm 0.68$ |
| | 0.25 | $3.47 \pm 0.75$ | $9.83 \pm 0.93$ | $19.20 \pm 1.10$ | $29.40 \pm 0.91$ | $39.72 \pm 0.91$ |
| | 0.5 | $4.40 \pm 0.83$ | $11.23 \pm 1.45$ | $23.49 \pm 1.02$ | $34.76 \pm 0.86$ | $45.98 \pm 1.09$ |
| | 0.75 | $2.36 \pm 0.84$ | $5.46 \pm 0.99$ | $10.47 \pm 0.99$ | $15.97 \pm 0.85$ | $21.10 \pm 1.10$ |
| | 1.0 | $-3.50 \pm 0.88$ | $-8.25 \pm 0.85$ | $-17.19 \pm 0.87$ | $-25.45 \pm 1.00$ | $-34.29 \pm 1.05$ |

Table S2: Simulated CLiP results for cohorts varying by total variance explained ($h^2$), cohort size, and percentage of individuals in the cohort that are true cases, with the remaining individuals being simulated controls. All trials were run with 100 SNPs with a fixed uniform effect size and an allele frequency of 0.2. Shown are mean and standard deviations of 20 trials.

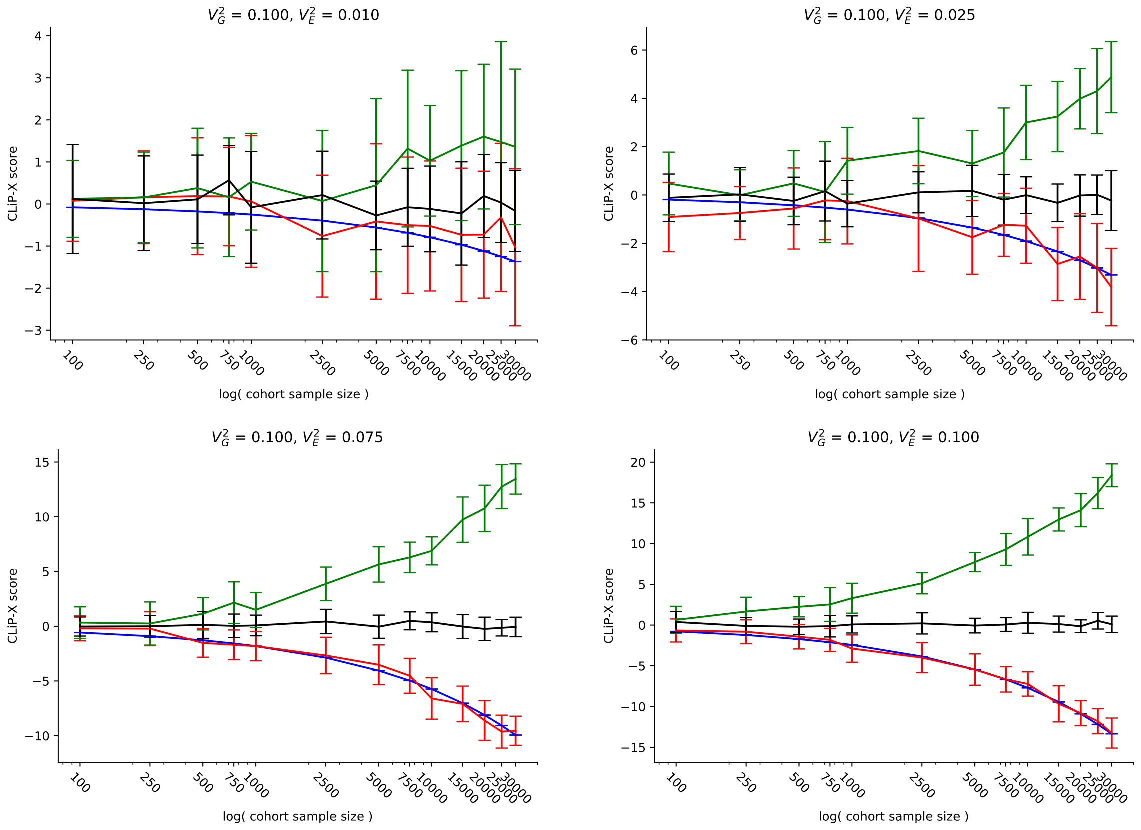


Figure S3: Additional heterogeneity scores for controls **(Black)**, homogeneous cases **(Red)**, and heterogeneous cases **(Green)** across different levels of variance explained by generated predictors ($V_E^2$). The predicted score for homogeneous cases **(Blue)**, calculated from summary statistics used to generate predictors from sampled SNPs and phenotypes from predictors, is the true null hypothesis of the heterogeneity test. Means and standard errors are shown for 20 trials over 10 expression variables generated by 100 SNPs.

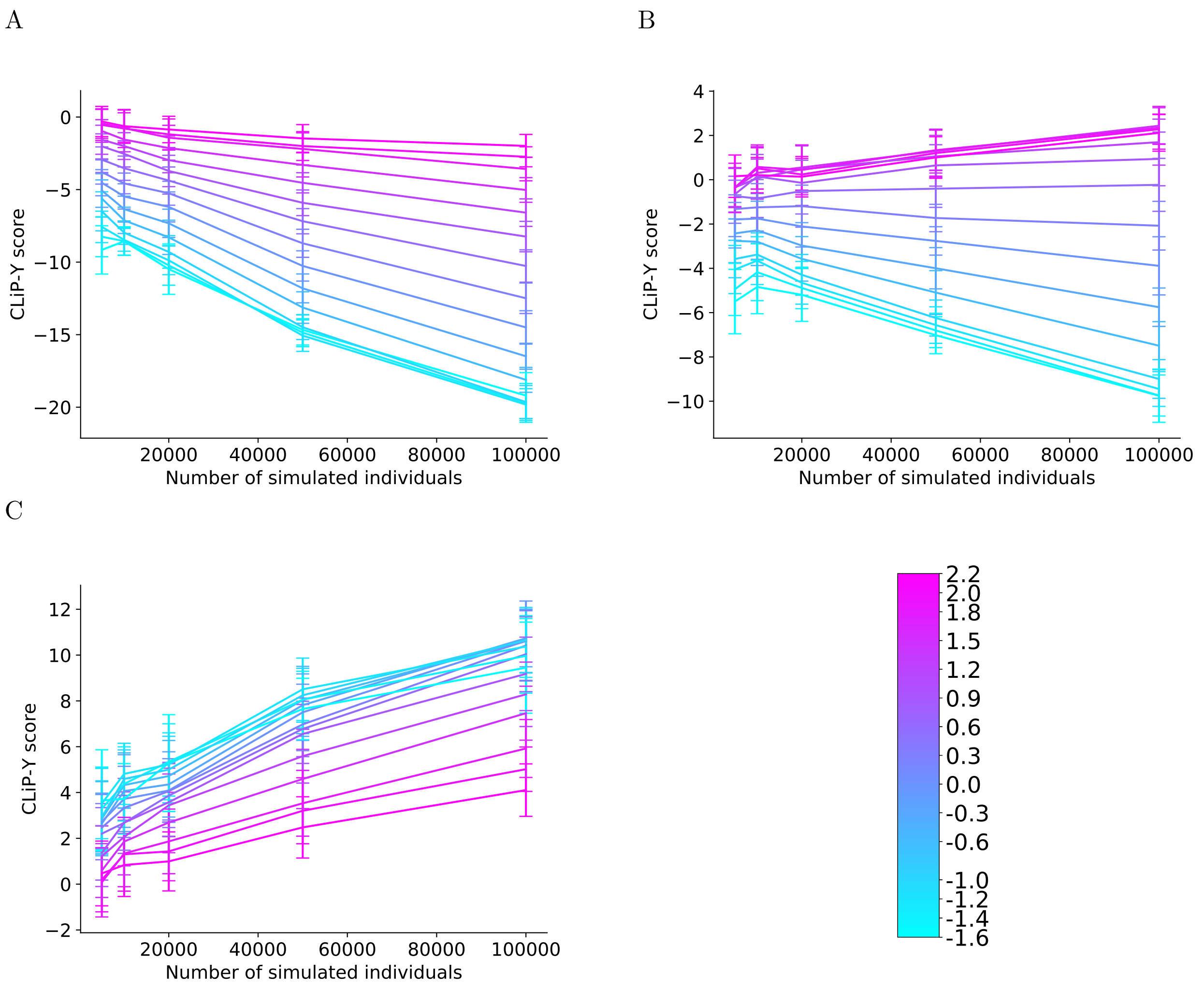


Figure S4: Heterogeneity scores for quantitative phenotypes split into artificial cases and controls by a hard threshold. Means and standard deviations of scores are shown as a function of sample size: **(A)** homogeneous cohorts, **(B)** heterogeneous cohorts, and **(C)** the difference (heterogeneous minus homogeneous) scores. The color gradient indicates the location of the threshold separating cases and controls. Each condition was run for 20 trials, and all cohorts were simulated with 100 SNPs and a total variance explained of 0.1.

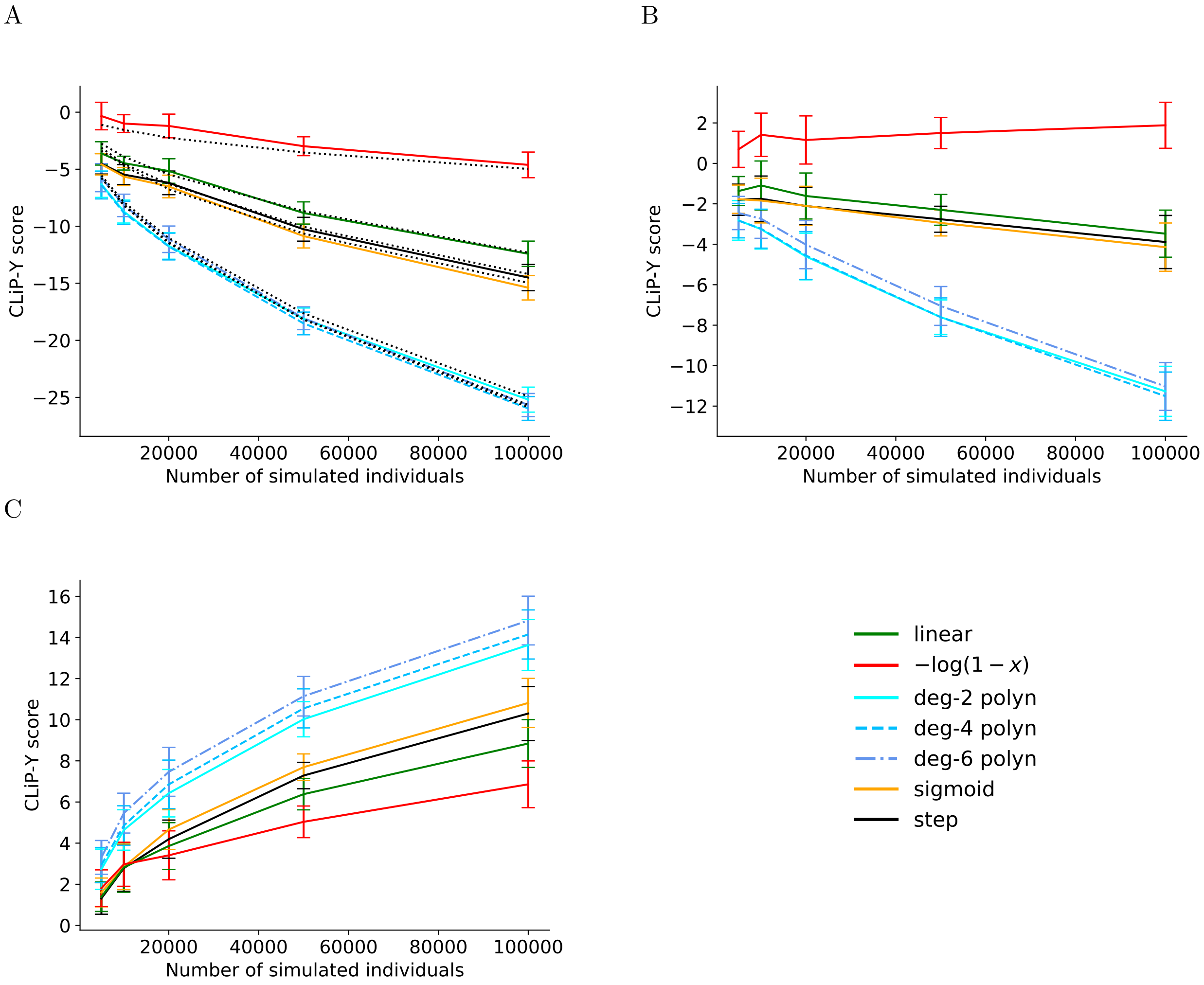


Figure S5: Heterogeneity scores for quantitative phenotypes with weight functions over individuals contributing to the correlation. Means and standard deviations of scores are shown as a function of sample size: **(A)** homogeneous cohorts, **(B)** heterogeneous cohorts, and **(C)** the difference between scores of heterogeneous cohorts and expected homogeneous scores in A. Colors indicate the type of weight function, with blue lines indicating learned polynomial functions. In **(A)**, black dotted lines indicate calculated expected scores for each homogeneous cohort, given effect sizes and allele frequencies. In practice, these expected scores will serve as the null scores subtracted from that of test data to produce analogous results to **(C)**. Each condition was run for 20 trials, and all cohorts were simulated with 100 SNPs and a total variance explained of 0.1.

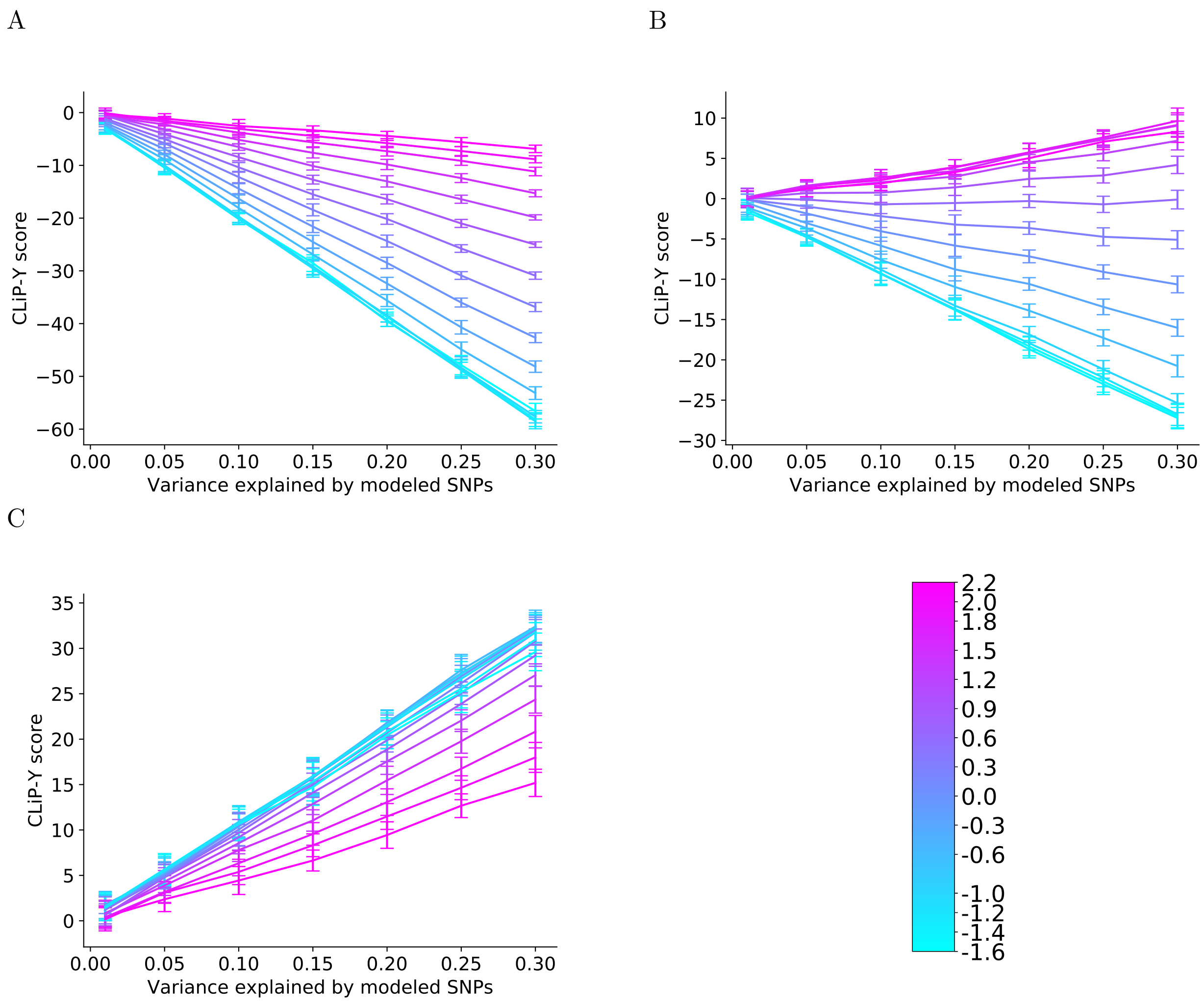


Figure S6: Heterogeneity scores for quantitative phenotypes with weight functions over individuals contributing to the correlation. Means and standard deviations of scores are shown as a function of total variance explained by SNPs: **(A)** homogeneous cohorts, **(B)** heterogeneous cohorts, and **(C)** the difference between scores of heterogeneous cohorts and expected homogeneous scores in A. Colors indicate the type of weight function, with blue lines indicating learned polynomial functions. In **(A)**, black dotted lines indicate calculated expected scores for each homogeneous cohort, given effect sizes and allele frequencies. In practice, these expected scores will serve as the null scores subtracted from that of test data to produce analogous results to **(C)**. Each condition was run for 20 trials, and all cohorts were simulated with 100 SNPs and a sample size of 100,000.

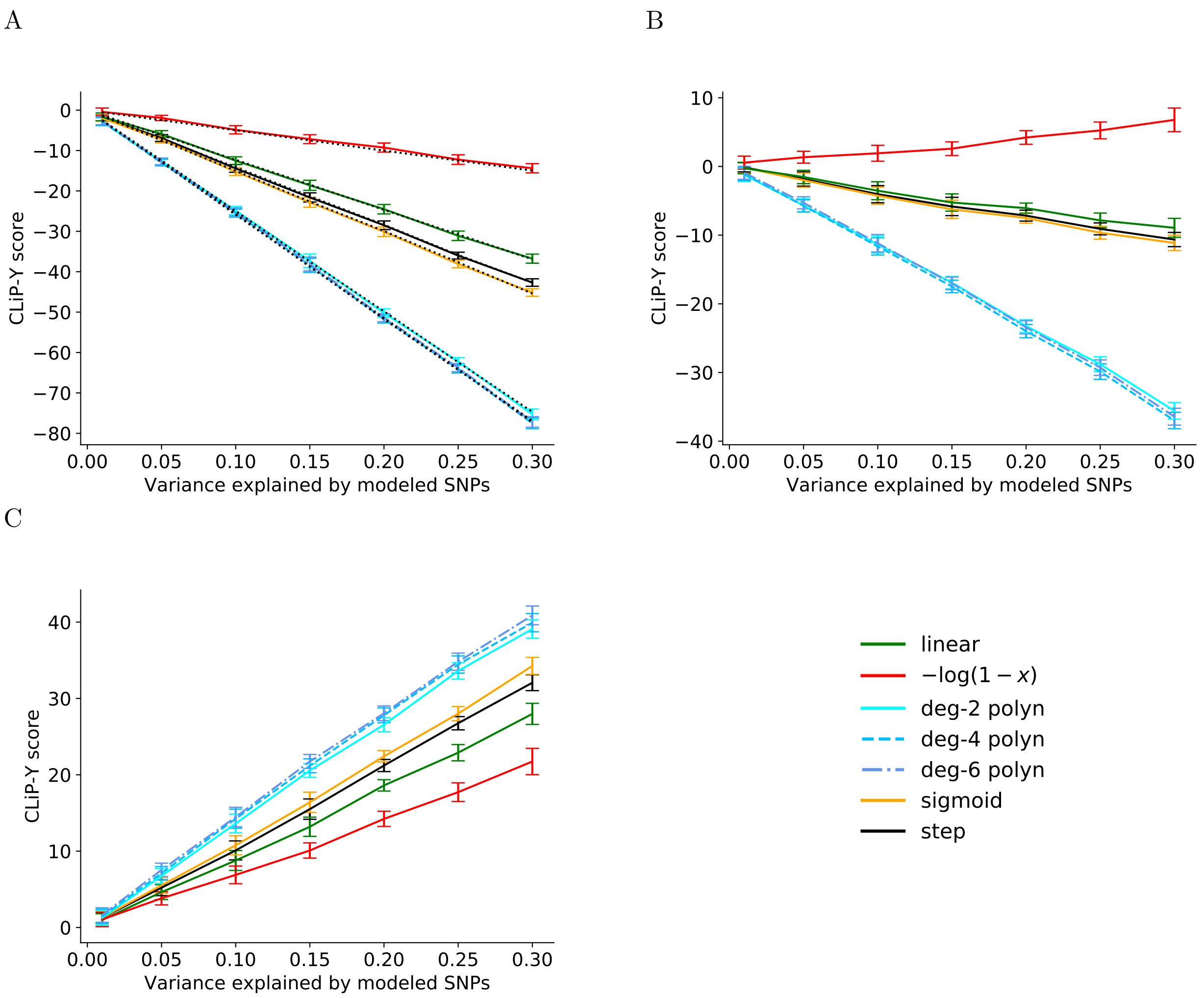


Figure S7: Heterogeneity scores for quantitative phenotypes with weight functions over individuals contributing to the correlation. Means and standard deviations of scores are shown as a function of variance explained: **(A)** homogeneous cohorts, **(B)** heterogeneous cohorts, and **(C)** the difference between scores of heterogeneous cohorts and expected homogeneous scores in A. Colors indicate the type of weight function, with blue lines indicating learned polynomial functions. In **(A)**, black dotted lines indicate calculated expected scores for each homogeneous cohort, given effect sizes and allele frequencies. In practice, these expected scores will serve as the null scores subtracted from that of test data to produce analogous results to **(C)**. Each condition was run for 20 trials, and all cohorts were simulated with 100 SNPs and a sample size of 100,000.

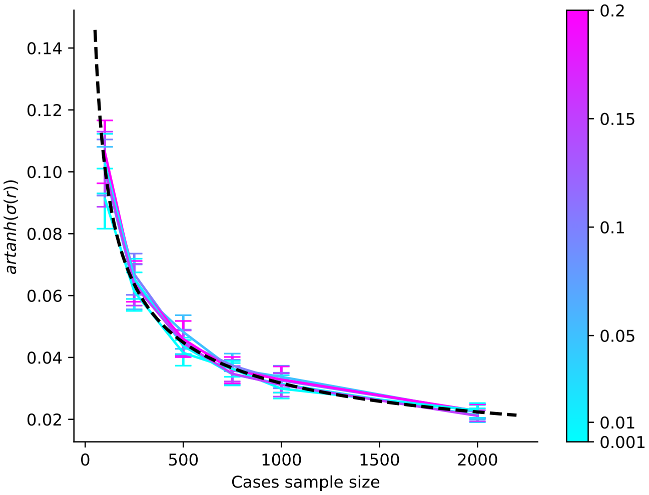


Figure S8: Validation of expected correlations in CLiP-X between expression variables in cases, accounting for contributions from PRS thresholding and shared SNP-expression effects. Shown are the outputs of an inverse hyperbolic function (artanh) of average standard deviations between predicted and generated correlations as a function of the case sample size used to estimate the correlation. Standard deviations across all pairs of variables are averaged for each experiment. The black line denotes the function $\frac{1}{\sqrt{N-3}}$, the expected standard deviation of the Fisher transformation for sample correlations.

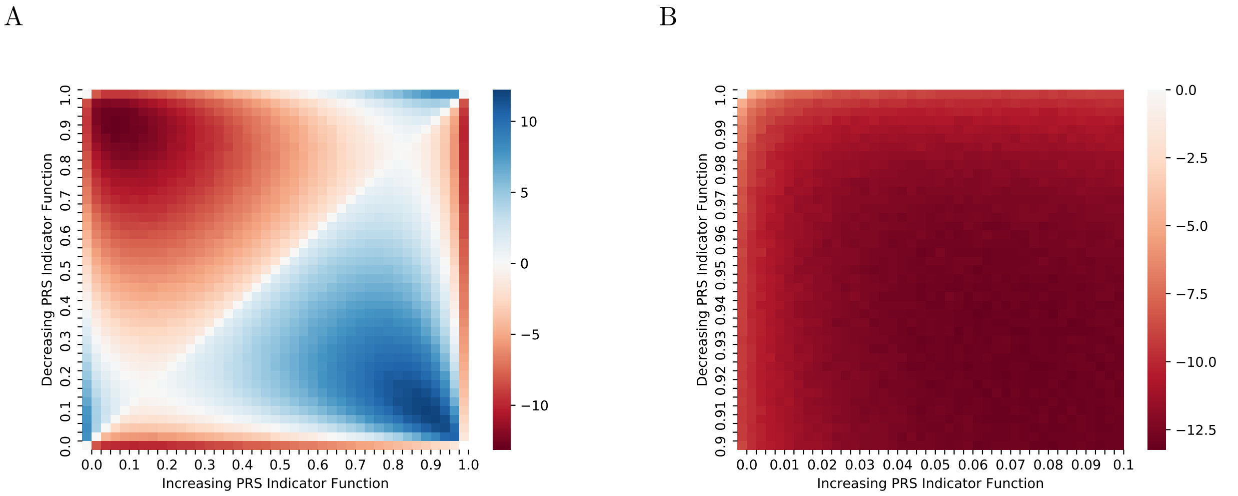


Figure S9: To demonstrate that optimal quantitative weight functions for heterogeneity are concave functions, two interval indicator functions in $[0, 1]$, an increasing one for $[x, 1]$ (x-axis) and a decreasing one for $[0, y]$ (y-axis) are combined so that their sum is the tested weight function. Each bin on the axes represents a transition point for the two step functions. The heterogeneity score is tested against a single homogeneous cohort, so optimal scores should be those that are most negative. **(A)** The best scores are those where $x$ is low on the PRS percentile scale but not 0, while $y$ is high on the PRS percentile scale but not 1. This coincides with the optimal polynomial functions obtained by a local search. **(B)** A zoomed view of the green box in A, showing that the optimal scores are not obtained by step functions at the periphery of the PRS distribution.
